# Mentalizing in an economic games context is associated with enhanced activation and connectivity in left temporoparietal junction

**DOI:** 10.1101/2022.02.12.480201

**Authors:** Li-Ang Chang, Lotte Warns, Konstantinos Armaos, Ava Q. Ma de Sousa, Femke Paauwe, Christin Scholz, Jan B. Engelmann

## Abstract

Studies in Social Neuroeconomics have consistently reported activation in social cognition regions during interactive economic games suggesting mentalizing during economic choice. It remains important to test the involvement of neural activity associated with mentalizing in an economic games context within the same sample of participants performing the same task. We designed a novel version of the classic false-belief task in which participants observed interactions between agents in the ultimatum and trust games and were subsequently asked to infer the agents’ beliefs. We compared activation patterns during the economic-games false-belief task to those during the classic false-belief task using conjunction analyses. We find significant overlap in left TPJ, and dmPFC, as well as temporal pole during two task phases: belief formation and belief inference. Moreover, gPPI analyses show that during belief formation right TPJ is a target of both left TPJ and right temporal pole (TP) seed regions, while during belief inferences all seed regions show interconnectivity with each other. These results indicate that across different task types and phases, mentalizing is associated with activation and connectivity across central nodes of the social cognition network. Importantly, this is the case in the context of the novel economic-games and classic false-belief tasks.

## Introduction

Inferring others’ mental states and predicting their intentions and beliefs is a social cognitive ability that supports social interactions. This ability is commonly referred to as “theory of mind” or “mentalizing”. Studies in social neuroscience have gathered substantial amounts of data on the neural networks involved in inferring others’ beliefs and intentions, yielding sufficient data to conduct meta-analyses with well over one hundred studies that jointly have identified consistent activations in a specific brain network (e.g., Mar, 2011; Molenberghs et al., 2016; Schurz et al., 2014, Decety and Lamm, 2007; Mitchel, 2009; Amodio and Frith, 2006, van Overwalle, 2009). The core mentalizing network identified by these studies consists of bilateral temporoparietal junction (TPJ), medial Prefrontal Cortex (mPFC), superior temporal sulcus (STS), temporal pole (TP) and precuneus (sometimes including posterior cingulate cortex, PCC).

Social neuroeconomics is another strand of research that has progressed relatively independently and that has repeatedly identified similar activation patterns within a similar network of brain regions when participants decide whether to cooperate with strangers in the context of economic games (for meta-analyses see Belluci et al., 2017; Feng et al., 2015; Schurz et al., 2014). The striking overlap of activations when participants perform classic false-belief tasks designed to study basic mentalizing processes, and when they make decisions in the context of economic games (see **Figure S1** for a neurosynth meta-analysis results that show this overlap) has been taken to suggest that participants engage in belief-based inferences that rely on mentalizing about their interaction partners when making interactive economic decisions (Fehr and Camerer, 2007; Engelmann et al., 2019; Alos-Ferrer and Farolfi, 2019). Neuroimaging studies have consistently revealed such social cognitive activations during social decision-making in the context of the trust game (e.g., McCabe et al. 2001, Krueger et al., 2007; Engelmann et al, 2019; Krueger, Grafman, & McCabe, 2008; Sripada et al., 2009; Stanley et al., 2012). Similar social cognitive activations have also been observed during the ultimatum and prisoners dilemma games (for a detailed description of these games see Engemann, Bzdock, Eickhoff, Vogeley, Schilbach, 2012). Results from an initial study on the neural correlates of trust decisions demonstrated activation of the dmPFC during social vs. non-social interactions in cooperative players (McCabe et al. 2001). This involvement of social cognition regions during trust decisions has been replicated and extended in subsequent studies, which also show recruitment of a wider social cognition network that includes dmPFC, TPJ and STS across different experimental contexts (Krueger et al., 2007; Engelmann et al, 2019; Krueger, Grafman, & McCabe, 2008; Sripada et al., 2009; Stanley et al., 2012). In fact, a recent study identified a wider network of regions consisting of dmPFC, Anterior Insula (AI) and pSTS that is more strongly interconnected with left temporoparietal junction during trust decisions and in people that are more trusting on average (Engelmann et al., 2019). The trends reflected in these findings are confirmed by a recent meta-analysis by Feng et al. (2015) that shows activations in precuneus, dmPFC and STS when participants consider unfair (relative to fair) offers.

The notion that the activation of social cognition regions during interactive economic games reflects mentalizing is further supported by theoretical considerations (Rilling and Sanfey, 2011; Engelmann et al., 2019; Fehr and Camerer, 2007; Alos-Ferrer and Farolfi, 2019). In economic games, mutual cooperation typically leads to financial gains for both interaction partners. However, there is a flip side in which financial losses can occur if one interaction partner decides to act selfishly to obtain higher payouts for herself at the cost of the other (Engelmann and Fehr, 2017). Because strategic interactions involve this possibility of non-cooperation by one of the interaction partners, participants have a strong incentive to avoid such outcomes of losing their initial investment (Aimone, Houser, & Weber, 2014; Bohnet & Zeckhauser, 2004; Bohnet, Greig, Herrmann, & Zeckhauser, 2008). In experimental games, the best way to assess the likelihood of non-cooperation is by taking the perspective of the interaction partner, i.e., mentalizing, which allows the participant to simulate how an interaction partner might act given the rules of the game. Activations in social cognition regions at the time point at which participants decide whether to invest an amount of money into another person therefore likely reflect mentalizing that aims to assess the degree of strategic uncertainty in a given context. This notion is further supported by results from repeated economics games, in which participants learn about the trustworthiness of interaction partners over the course of multiple trials. Feedback about partners’ decisions activates social cognition regions in dmPFC, TPJ and PCC (Rilling et al., 2004). Similarly, when building trust during the early stages of a repeated interaction dmPFC is active, while it is relatively deactivated once trust has been established in the later stages of repeated games (Krueger et al., 2007). In fact, learning about the characteristics of interaction partners has repeatedly been associated with prediction error signals in social cognition regions (Behrens et al., 2008). These results directly implicate social cognitive processes computed in this network in learning and updating about the prosocial characteristics of current interaction partners.

Despite these theoretical considerations and the considerable overlap of activations in the mentalizing network during false-belief tasks and economic games, one major shortcoming of this strand of research is that evidence for evoking mentalizing and social cognitive processes during social decision-making within economic games is indirect and to date relies largely on reverse inference (Poldrack, 2006). To directly test the involvement of mentalizing processes during interactive economic decisionmaking, it is necessary to explicitly assess participants’ thoughts about the mental states of others within the context of economic games.

To this end, we combine the approaches developed by the two research streams of social neuroscience and social neuroeconomics. Specifically, we developed a novel false-belief task (FBT) that required participants to apply the rules of two well-established economic games, the trust and ultimatum game, to be able to correctly answer incentivized questions that assessed our participants’ understanding of economic game interactions. In our novel economic game version of the FBT, participants first read about an interaction between two parties and were then asked to either infer the false belief of one of the interaction partners in the belief condition, or to calculate the payoff for one of the interaction partners in the outcome condition, which does not require mentalizing. The false belief condition therefore assessed our participants’ understanding of how different economic game situations might cause false beliefs held by one of the interaction partners, while the outcome condition allowed us to assess our participants’ understanding of the rules of the game and how payouts were computed. Our approach therefore enabled us to directly assess belief-based inferences in the context of economic games and compare the activation patterns during belief-based inferences in the context of economic games to those during the standard false-belief task. Of note, using this approach in which our participants form beliefs about others based on their observations of two agents that interact in the context of economic games has the distinct advantage that their mentalizing processes are not distorted by additional cognitive and affective processes that occur in direct interactions within economic games and the decision processes associated with that. Our approach therefore controls for the distortionary influences of valuation processes, strategic considerations, reputation concerns, fairness considerations and other social preferences, as well as affective reactions that are common to direct trust and ultimatum game interactions, thereby allowing us to cleanly disentangle mentalizing processes in the context of economic games.

Given the strong suggestion from theoretical considerations, prior research and metaanalyses, we expected that belief-based inferences (relative to outcome-based inferences) in the context of economic games lead to similar activation patterns within the mentalizing network as the standard false-belief task. Moreover, given similar average activation patterns, activity within key regions may also be similarly interconnected across the two contexts. Thus, we also assessed the functional connectivity of the mentalizing network averaged across the two task contexts.

## Materials and Methods

### Participants

Two pilot studies were conducted to develop and further titrate the novel game-theoretic vignettes. Pilot experiment 1 was conducted online via Qualtrics with 50 participants (33 females, age mean = 33.4 years, SD = 8 years) that were recruited via Prolific. Pilot experiment 2 was conducted at the Center for Research in Experimental Economics and Political Decision Making (CREED) with 38 participants (26 females, age mean = 21.9 years, SD = 1.9 years). All procedures for pilot experiments were approved by the ethics committee of economics and business at the University of Amsterdam.

39 right-handed volunteers participated in the main fMRI experiment (18 males, aged 18 - 33, mean (SD) = 22.51 (4.03) years) mainly recruited from the participant pool of the Behavioural Science Lab of the Faculty of Social and Behavioral Sciences at the University of Amsterdam (LAB, https://www.lab.uva.nl/lab/home). All participants first underwent an initial screening, which required that participants (1) were between 18 and 40, 2) were right handed, 3) had no history of any neurological or mental illness, 4) were fluent in English, 5) agreed to receiving mild electric shocks during the experiment, 6) never participated in a corresponding behavioral pilot study previously conducted as part of this study, and 7) fulfilled all MRI-safety requirements according to the guidelines of Spinoza Center of the University of Amsterdam. Two participants were excluded from further analysis due to excessive head movement (>2 x voxel size (6 mm), 1 participant), and due to low accuracy of responses (mean accuracy < 3 (SD) of sample mean, 1 participant). The final dataset for fMRI analysis therefore consisted of 37 participants. Written informed consent was obtained from all participants before their participation. All procedures were implemented in compliance with the guidelines formulated by the Ethics Review Board of the Faculty of Social and Behavioral Sciences, University of Amsterdam.

### Pilot experiments

We first developed a set of game-theoretic vignettes by outlining a number of interaction scenarios from economic games that reflect false beliefs of one of the interaction partners. In these scenarios, we built upon two well-established economic games, the trust game and the ultimatum game, which can be easily explained to participants (see the *stimuli section* for a detailed description of the novel scenarios, and our project page on osf.io for detailed instructions https://osf.io/3eg56/?view_only=face48878dd144848d26f1c7d3c47d31). Aim of an initial pilot study that was conducted online via Prolific was to test participants’ understanding of the different vignettes, and to identify potential outlier scenarios that might not be easily understood by our participants. Vignettes and subsequent questions that probed participants understanding were presented to participants via Qualtrics, and reaction times were recorded. The response times indicated that trust game outcome vignettes were perceived as too difficult among the four conditions included in this pilot study [TG outcome average RT =24.68s, se=1.69, UG outcome average mean RT = 15.80s, se=0.84, TG belief average RT=15.13s, se=1.00, UG belief average RT=17.17, se=0.89]. Because paired t-tests showed significantly longer RTs in the TG outcome condition compared to all other conditions (UG outcome, t(49)=6.79, *p*=1.76×10^−8^; TG belief, t(49)=6.37, *p*=6.24×10^−8^; UG belief, t(49)=4.82, *p*=1.43×10^−5^), we simplified the computations required for correct responses by restricting possible answers to multiples of five in the trust game outcome scenarios.

Next, we validated our new stimulus set in the laboratory by conducting an additional behavioral pilot conducted in the CREED laboratory. This experiment allowed further fine-tuning of the final set of vignettes and experimental parameters such as the appropriate difficulty and timing of stimuli. The experimental design was equivalent to the design reported for the fMRI experiment below, except that participants were also required to indicate when they completed reading during the vignette period by pressing the space bar. While participants were reminded of this in the instructions, we received a relatively low response rate (32% of all trials) indicating that participants had difficulties with the dual task of reading and button pressing within the given period of time. Given these difficulties and to allow participants to fully concentrate on reading the vignettes and to avoid confusion during the fMRI experiment, no button presses were required during the vignette period in the fMRI experiment. Participants were paid on a piece-rate basis (20c per correct answer) and received an average of 28.38 Euros for their participation (average piece rate earnings of 18.38 plus 10 Euros for completing the online survey). Accuracy and response time results from the pilot study are reported alongside results from the main fMRI experiment in tables 1 and 2.

**Table 1.**
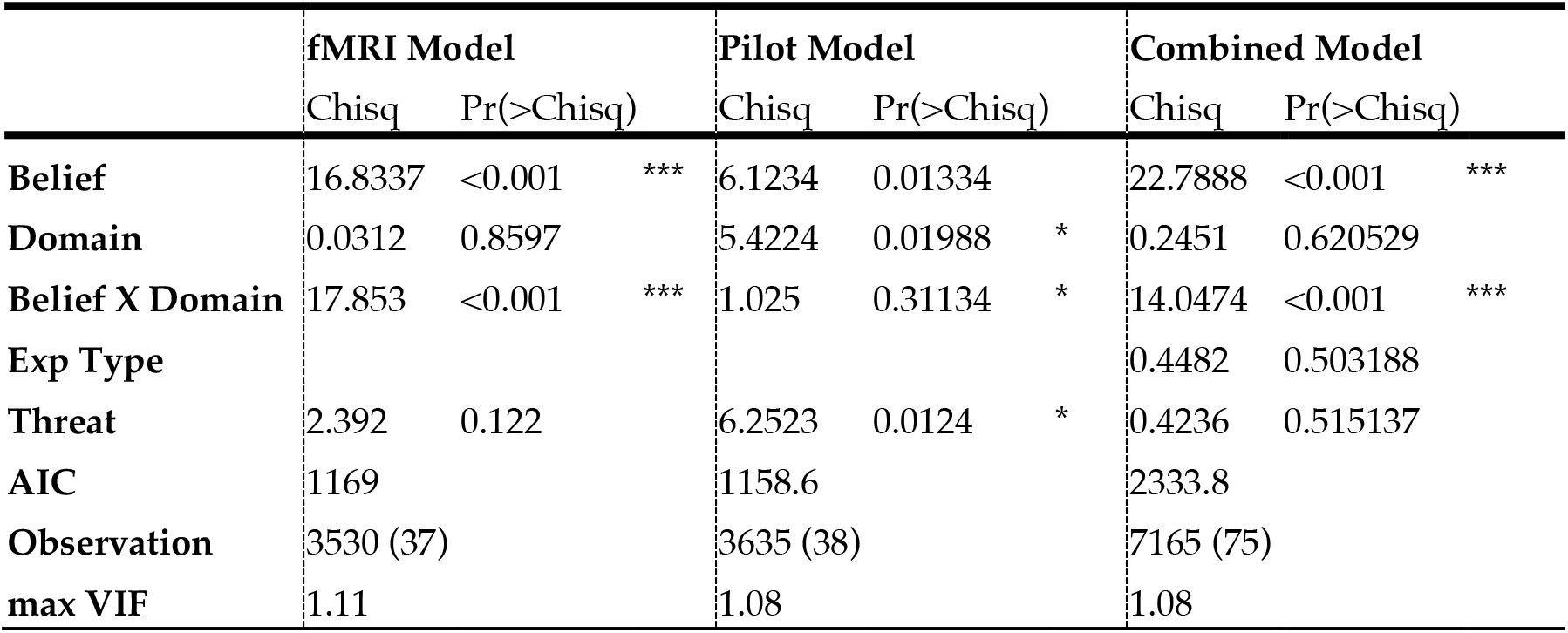
ANOVA tables for accuracy for three different models that include the fMRI, pilot and combined datasets. Models use a restricted random effects structure with random slopes for the Task Domain factor (except for the pilot model) and random intercepts and were estimated using the AFEX package. ANOVA results are based on logistic regressions with correct/incorrect responses as dependent variable.

**Table 2.**
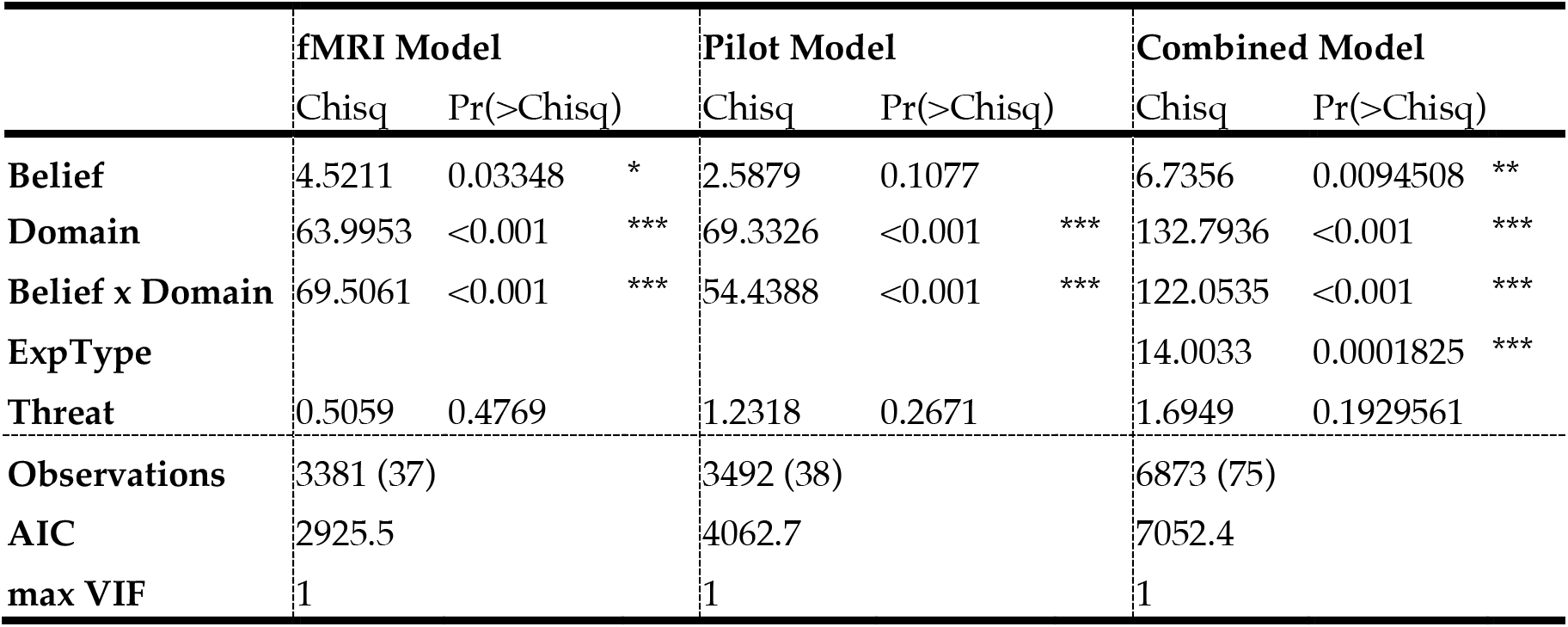
ANOVA tables for log RT for three different models that include the fMRI, pilot and combined datasets. All models use a maximal random effects structure with random slopes and intercepts and were estimated using the AFEX package. Dependent variable is the logarithm of RT for correct trials only.

|  | fMRI Model |  |  | Pilot Model |  |  | Combined Model |  |  |
| --- | --- | --- | --- | --- | --- | --- | --- | --- | --- |
|  | Chisq | Pr(>Chisq) |  | Chisq | Pr(>Chisq) |  | Chisq | Pr(>Chisq) |  |
| **Belief** | 4.5211 | 0.03348 | * | 2.5879 | 0.1077 |  | 6.7356 | 0.0094508 | ** |
| **Domain** | 63.9953 | <0.001 | *** | 69.3326 | <0.001 | *** | 132.7936 | <0.001 | *** |
| **Belief x Domain** | 69.5061 | <0.001 | *** | 54.4388 | <0.001 | *** | 122.0535 | <0.001 | *** |
| **ExpType** |  |  |  |  |  |  | 14.0033 | 0.0001825 | *** |
| **Threat** | 0.5059 | 0.4769 |  | 1.2318 | 0.2671 |  | 1.6949 | 0.1929561 |  |
| **Observations** | 3381 (37) |  |  | 3492 (38) |  |  | 6873 (75) |  |  |
| **AIC** | 2925.5 |  |  | 4062.7 |  |  | 7052.4 |  |  |
| **max VIF** | 1 |  |  | 1 |  |  | 1 |  |  |

### fMRI experiment

#### fMRI Experiment Procedure

Participants were first invited to complete an online prescreening questionnaire and a battery of personality measures via Qualtrics before the main fMRI experiment. Participants were given 14 Euros for completing this online survey. In part two of the experiment, participants were invited to the fMRI laboratory at the Behavioral Science Lab of the University of Amsterdam. They were asked to read the instructions thoroughly and complete a quiz afterwards to ensure they fully understood the task, especially the rationale behind the economic games (Trust Game, TG; and Ultimatum Game, UG, for instructions see our project page on osf.io https://osf.io/3eg56/?view_only=face48878dd144848d26f1c7d3c47d31). They were allowed to ask questions during the instructions and the quiz, which the experimenters answered carefully. In addition, participants had the opportunity to practice the task before the fMRI experiment. To further ensure participants’ comprehension of the task, all participants were required to achieve at least 66% accuracy before proceeding to the main experiment. Among all participants, only three required two practice runs, after which they passed the threshold of correct answers. After being placed in the scanner, participants underwent a short button training task to allow familiarization with the button box. Subsequently, they completed four fMRI runs, with each run consisting of 24 trials that were subdivided into 8 blocks of 3 trials each. Participants also underwent electrical stimulation calibration before the 1^st^ and 3^rd^ run (for details see Engelmann et al., 2019) to determine pain thresholds for the Threat condition, which we control for, but do not specifically analyze in the current set of analyses. After scanning, participants filled out an exit questionnaire, after which they were paid their show-up fee and performance bonus.

##### Vignette Stimuli

A novel set of vignettes was developed for the current study, with the aim to test the neural correlates of belief formation and inferences in the context of economic games. These were combined with vignettes from prior research (Saxe & Kanwisher, 2003; Bruneau, Pluta & Saxe, 2012), to enable comparisons with the well-established false-belief task. The novel economic game vignettes described interactions between agents in the trust and ultimatum games and therefore required an understanding of the rules of these games, which were explained in detailed instructions. Economic game scenarios were based on 6 different hypothetical events that can occur in laboratory contexts. Importantly, in all scenarios one interaction partner keeps all, or the majority of the accumulated money for different reasons. The reasons included the participant’s decision to invest their winnings into charity, and incorrect decisions due to a computer error, accidentally pressing the wrong button, or because of misunderstanding the game setup. Example vignettes are shown in Figure 1A, and the complete list of economic game vignettes can be found on our project page on osf.io (https://osf.io/3eg56/?view_only=face48878dd144848d26f1c7d3c47d31). Additionally, two types of questions were developed that probed participants’ understanding of the interactions described in the vignettes: one type focused on the false belief of one of the agents, while the other type focused on the payouts for one of the agents.

**Figure 1.**
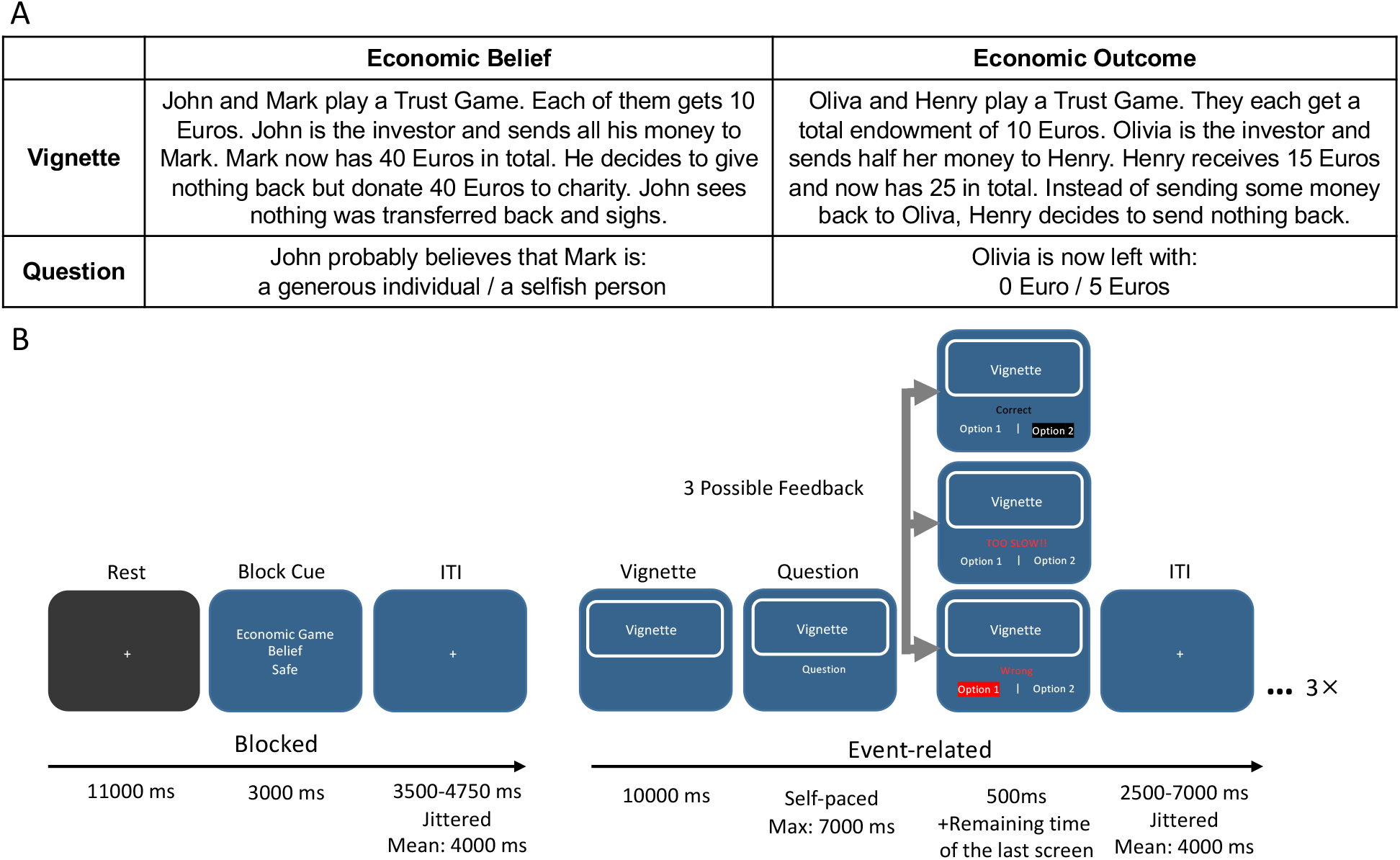
Example Economic Game Vignettes and Task Schematic. (A) A set of novel vignettes based on economic games were developed for the current experiment. The examples in A show economic game vignettes in the Belief (left) and the Outcome (right) condition, together with their respective questions. (B) Trial sequence of fMRI experiment. An initial block cue indicated the conditions that remained stable for the duration of one block of three trials, including the domain of the vignette, and whether the vignette concerns beliefs or outcomes. The vignette (see A) was shown for 10 seconds, after which participants were given a maximum of 7 seconds to answer the question. Correct answers were incentivized with a piece rate of 0.2 Euro.

Given the novelty of the task, we also assessed whether our participants used a strategy to answer questions about economic game interactions in an open-ended question that was included in the exit questionnaire. We find that a subset of participants indeed used a strategy to answer questions about economic-game vignettes. We therefore reanalyzed our behavioral and imaging data by including a binary covariate for strategy in our behavioral and fMRI models (reported in Tables S1-3). Our results indicate that there were no significant modulatory effects of using a strategy on the behavioral and imaging results of the economic-games vignettes.

A total of 4 different vignette types were included in the experiment, and varied along the experimental factors Domain (life stories vs. economic games) and Belief (false belief vs. outcome description). Note that participants also performed half the trials under Threat induced through a probabilistic electric shock (threat present vs. threat absent), which in the current analyses we control for, but do not specifically analyze (see Chang et al., in prep., for this analysis). Specifically, in the Life Story-Outcome condition, the participants were reading about events that happen to another person. They were asked to answer questions about an objective description of the consequence of the event. In the Life Story-Belief condition, the participants were explicitly asked about the most likely beliefs or intentions of the protagonist in the scenario. On the other hand, in the Economic Game-Outcome condition, the participants were asked to calculate the payoff of one of the interaction partners based on the rules of the economic game in question (TG or UG). Note that this condition served not only as a contrast condition in the economic game domain, but also allowed us to probe our participants understanding of the rules of the economic games reflected by (in)correct calculations of the payouts across different game contexts. Similar to the belief condition in the life story domain, in the Economic Game-Belief condition, the participants were required to infer the (false) beliefs of the interaction partners during an economic game. Figure 1A shows example economic game vignettes (for the full list of economic game vignettes, see our project page on osf.io https://osf.io/3eg56/?view_only=face48878dd144848d26f1c7d3c47d31).

In addition, the vignettes based on the trust game and ultimatum game were never presented together in the same block to avoid potential confusion and task switching. Furthermore, the different scenario types (i.e., life-belief, life-outcome, econ-belief, econ-outcome), game types (TG, UG) and scenario topics (e.g., computer error scenarios) were pseudorandomly distributed across Threat conditions.

##### Task description

Figure 1B illustrates the sequence and timing of a representative block and trial. Each block started with a block cue informing participants of the condition throughout the current block (3000 ms). Conditions varied based on the factors Task Domain (life story vs. economic games), Belief (false belief vs. outcome description), and Threat (threat present vs. threat absent), and were randomized throughout the experiment and for each participant. The example in Figure 1B shows an Econ-Belief-NoThreat condition, indicating that the 3 vignettes in the current block contain economic game scenarios, in which participants were asked to infer the interaction partner’s intentions and beliefs, and they did not receive electric shocks throughout this block. The block cue was then followed by a blank screen containing a fixation cross for a jittered duration (range: 3500ms – 4750ms, mean: 4000ms). Thereafter, participants were asked to read the current vignette, for which they were given 10000ms. This period is referred to as the vignette period below, during which participants read about a sequence of events that enabled them to develop an understanding of the protagonist’s beliefs in the belief condition as illustrated in Figure 1A. The vignette display was followed by a question period which was self-paced and terminated after 7000ms. During this period participants were required to integrate the information gathered during the vignette period to answer incentivized questions about the beliefs of one of the protagonists in the belief condition, as illustrated in Figure 1A. Participants chose from two possible options, one incorrect and one correct one, with the position of the correct option randomized across trials. Note that correct answers were incentivized at a piece rate of 0.2 Euro to ensure that participants maintain attention and motivation throughout the experiment (Contreras-Huerta et al., 2020). It was therefore in the best interest of participants to answer correctly and within the 7000ms period, as otherwise they would forgo payment on that trial. Feedback was shown for 500ms as soon as the participant pressed the corresponding button of the option, or after the 7000ms period expired with no button press. Feedback indicated whether responses were correct, incorrect, or too slow. Note that participants were not able to move through the experiment faster by responding faster during the question period as the remainder of the question period was added to the feedback duration if RT < 7000ms. An additional jitter period (range: 25000 – 7000ms, mean: 4000ms) was added at the end of each trial before the next trial started. Given our use of a hybrid design, a rest period of 11000ms was added at the end of each block to allow the BOLD signal to return to baseline. Each participant completed a total of 96 trials distributed across 32 blocks and 4 runs. The task was programmed and presented in MATLAB 2017b using the Cogent toolbox (http://www.vislab.ucl.ac.uk/cogent.php). Task stimuli were projected on a screen at the scanner head and were visible to the participant via a mirror mounted onto the head coil.

#### Payment determination

Participants earned a € 0.20 bonus for each correct answer that was provided within the time limit of 7 seconds. The final payment for participation consisted of the performance bonus (max. 19.20 Euros) and the endowment of € 14 paid for completing the online survey before the fMRI experiment. Participants earned an average of € 32.32.

### Functional magnetic resonance imaging

#### FMRI data acquisition

fMRI data were collected using a 3.0 Tesla Philips Achieva scanner located at the Behavioral Science Lab at the University of Amsterdam. T1-weighted structural images were acquired (1 × 1 × 1 mm voxel size resolution of 220 slices, slice encoding direction: FH axial ascending, without the slice gap, TR = 8.2 ms, TE = 3.7 ms. flip angle = 8°). Functional images were acquired using a T2*-weighted gradient-echo, echo-planar pulse sequence (3.0 mm slice thickness, 3.0 × 3.0 mm in-plane resolution of 36 slices, slice encoding direction: FH axial ascending, slice gap = 0.3 mm, TR = 2000 ms, TE = 28 ms, Flip angle = 76.1°, and with 240 mm field of view). In addition, to correct EPIs for signal distortion, we also conducted an additional field-map scan at the half-way point of the experiment using a Phase-difference (B0) scan (2.0 × 2.0 × 2.0 mm voxel size resolution, axial ascending direction, without slice gap, TR = 11 ms, TE_s_ = 3 ms, TE_l_ = 8ms, flip angle = 8°).

#### FMRI preprocessing and analyses

Imaging data analysis was carried out with SPM12 (Wellcome Department of Cognitive Neurology, London, UK) and the CONN toolbox (Whitfield-Gabrieli & Nieto-Castanon, 2012). Preprocessing followed the following steps: First, all functional images were simultaneously realigned to the first volume of the first run using septic b-spline interpolation and unwarped (using B0 maps) using the realign and unwarp function in SPM, followed by slice timing correction. Afterward, T1-weighted structural images were co-registered with the functional images and then segmented into six different tissues classes using the segment function in SPM12. Next, all images were normalized to the Montreal Neurological Institute (MNI) T1 using the forward deformation parameters from segmentation. Lastly, all functional images were smoothed using spatial convolution with a Gaussian kernel of 6 mm at full width half maximum (FWHM).

Statistical analyses were carried out using the general linear model (GLM). To reflect our factorial design, the model included separate regressors of interest for each Domain (life story vs. economic games) and Belief (false belief vs. outcome description) condition for both the vignette reading period, and the question period. Regressors of interest were modeled using a canonical hemodynamic response function (HRF). To best capture mentalizing during the question period, we used a variable epoch model from the onset of the question until option choice (button press). We also modeled regressors of no interest, which include each block cue, the feedback period, shock moment and Threat condition (threat present vs. threat absent), as well as omitted trials in which no response was provided by the participant. While omissions were rare (on average 0.55%), these were modeled explicitly to ensure that we only included trials for which we are certain participants paid attention to the task. In addition, the six motion parameters derived from the realignment procedure were modeled as regressors of no interest. All results were FWE-corrected at cluster level with a cluster-forming threshold of *p* < 0.001.

Conjunction analyses were conducted to test the overlap between belief-based activations in the life story and the economic game domains and were based on the conjunction null (Nichols et al., 2005). Whole-brain statistical maps for each domain used a voxel threshold at an alpha value of *p* < 0.001 and were FWE corrected at the cluster level (for completeness we also report the uncorrected results in Table 5). The individual maps were then multiplied together using the ImCalc function in SPM12, which creates a map of voxels that are significantly activated in both conditions, reflecting a logical “and” conjunction (Nichols et al., 2005).

#### Connectivity Analyses

Generalized Psychophysiological Interaction (gPPI) analyses were conducted using the CONN functional connectivity toolbox (www.nitrc.org/projects/conn) (Whitfield-Gabrieli & Nieto-Castanon, 2012) using two-analysis approaches: (1) ROI-to-ROI analyses to identify the specific interconnectivity among a restricted set of regions of interest that are commonly associated with social cognitive processes, and (2) seed-based, whole-brain (seed-to-voxel) analysis to identify the wider connectivity of these social cognition regions with additional brain areas. Data were first prepared for connectivity analyses by preprocessing the fMRI data using the indirect segmentation and normalization pipeline in CONN, which is largely equivalent to our preprocessing steps above, but included the additional step of identifying and removing outlier scans from the analysis (ART, Whitfield Gabrieli). Next, the data underwent denoising. In accordance with the anatomical component-based noise correction method (aCompCor, Behzadi et al., 2007, Muschelli et al., 2014), denoising was conducted before functional connectivity analyses and included 10 CSF and 10 white matter principal components as nuisance covariates, as well as 6 realignment parameters, their first-order temporal derivatives and quadratic effects (24 parameters in total), the outlier scans identified by ART, and all task effects and their first-order derivatives (48 parameters in total). Low-frequency fluctuations were isolated using a low-pass temporal filter (.008 Hz) after denoising. Thresholding for ROI-to-ROI analyses was done using the Threshold-free cluster enhancement method (Smith and Nichols, 2007) with peak-level family wise error corrected p-values.

Seed regions for functional connectivity analyses were extracted from the conjunction maps assessing the activation overlap between economic-game and story-based task domains for vignette (see Figure 4) and question periods (see Figure 6). Note that because we focus on regions that were jointly activated during the economic-games and standard FBT and therefore have similar belief-based activation profiles, we did not distinguish between these task domains in connectivity analyses and analyzed belief-based connectivity (belief > outcome) independent of task domain. Furthermore, to ensure that current activations match social cognition regions from prior studies, these were further conjoined with the smoothed (FWHM kernel of 1mm) neurosynth map obtained via an association test for the term “mentalizing”. To remove smaller regions, we used a cluster threshold of k ≥ 25, which led to the following seed regions (maps with our seed regions can be found on our project page on osf.io https://osf.io/3eg56/?view_only=face48878dd144848d26f1c7d3c47d31).): 1) during the vignette period seed regions for connectivity analyses included dmPFC (6, 56, 23, k=46), left TPJ (−48, −58, 26, k=89), right TP (48, 2, −31, k=79), and right MTG (51, −28, −4, k = 26); during the question period, seed regions for connectivity analyses included left TPJ (−60, −61, 20, k=99), dmPFC (−6, 56, 26, k=55), left MTG (−54, −28, −4, k=112), and bilateral TP (left: −54, 5, −25, k=168, right: 48, −4, −37, k=178).

### Behavioral Results

The focus of our behavioral analyses was to test whether our novel economic game vignettes yielded behavior that is comparable to the standard life story vignettes in terms of overall accuracy and reaction times. At first glance there seem to be only small differences in accuracy and reaction times across the two domains, with average accuracy reaching 95% for both the life story domain and the economic game domain. Closer inspection, however, revealed differences between the domains that seem to be largely driven by differences between the Outcome conditions in the life-story and economic-game vignettes. This difference is likely due to the economic game outcome condition requiring computations of payouts, whereas standard vignettes require an understanding of the story line and the mental state of the protagonist.

To analyze the choice data, we conducted logistic regressions implemented in the context of a generalized linear mixed-effects model (GLME). Models included responses on each trial (correct/incorrect) and log reaction time as dependent variables, as well as Task Domain and Belief condition as fixed effects predictor variables, and Threat as a fixed effects control variable. Models were estimated via the mixed function of the AFEX package in R (Singmann, Bolker & Westfall, 2016) that relies on the lme4 package. We report results from models with the maximum possible random-effects structure (Barr et al., 2013). For reaction times, linear regressions using a full model structure with random slopes for the Task Domain and Belief factors, in addition to random intercepts were employed used. For accuracy, logistic regressions were employed. Including all random slopes led to overfitting, requiring us to reduce the number of random slopes, such that all final models include a subjectwise random intercept, and a subset include a random slope for the Task Domain factor. Note further, that we report analyses for the pilot experiment, the fMRI experiment and the combined dataset in all tables, but focus our discussion of the results on the data collected during the fMRI experiment.

### Accuracy across Belief Conditions and Task Domains

As reflected in Figure 2A, we find a significant main effect of Belief (X^2^ = 16.83, *p* < 0.001) on accuracy and a significant interaction between Belief and Task Domain (X^2^ = 17.85, *p* < 0.001). Follow-up tests of the interaction were conducted using the free method implemented via the multcomp package (Hothorn, Bretz & Westfall, 2008). Results from pairwise comparisons using the Sidak correction indicate that these effects are due to a significantly lower accuracy in the economic games compared to the life story task in the Outcome conditions (estimate = −0.82, Z = −2.60, *p* = 0.018), while only a near-significant difference between the economic games and life stimuli was observed in the Belief conditions (estimate = 0.75, Z = 1.91, *p* = 0.056).

**Figure 2.**
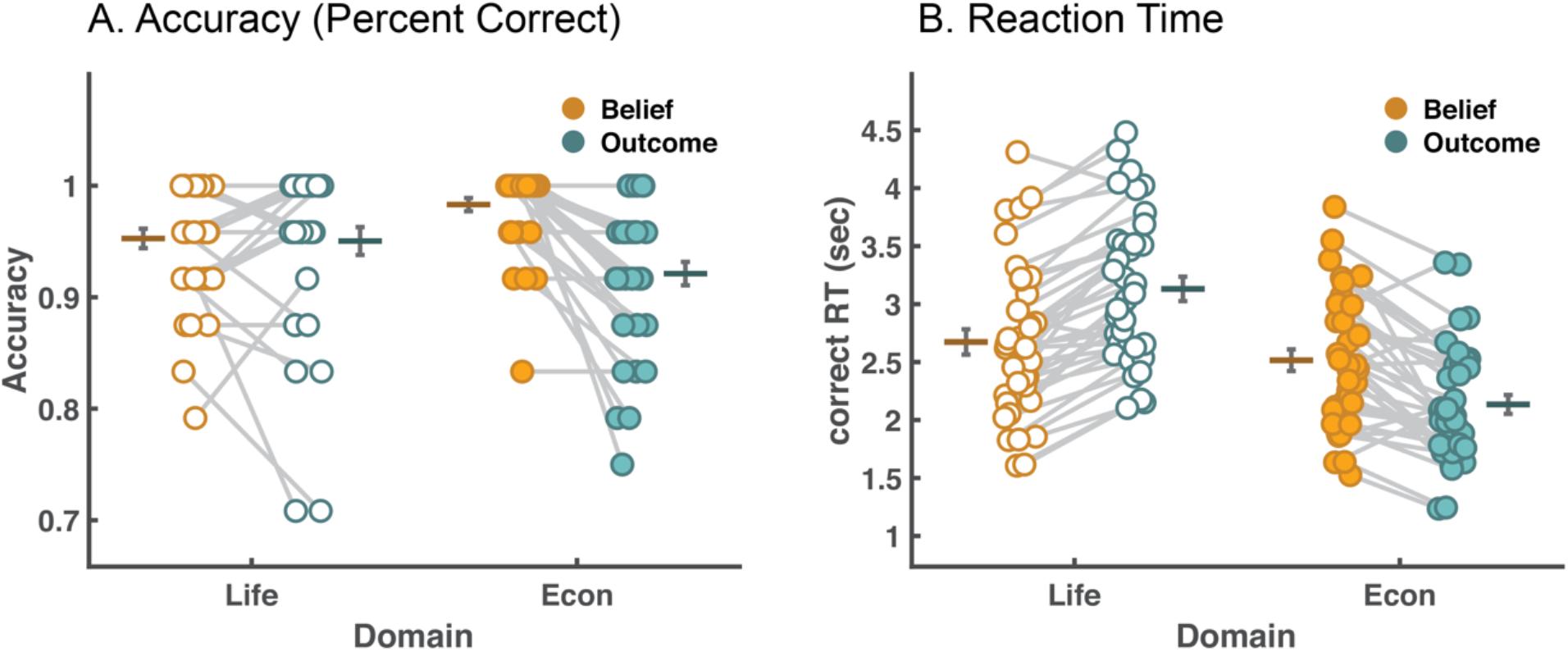
Behavioral Results. (A) shows the mean accuracy across (lines with standard error bounds) and within individuals (connected dots) of participants’ answers (percent correct) across Task Domain and Outcome conditions. Accuracy reflects the proportion of correct relative to all responses. (B) shows mean response times across (lines with standard error bounds) and within individuals (connected dots) for correct trials only across Task Domain and Outcome conditions.

This result indicates that accuracy differences were only found in the outcome, but not in the belief condition of the Belief Factor. The economic game outcome condition has different cognitive demands compared to those of all other conditions as it requires computations of payouts, which is reflected by the current results. Note that, except for the belief main effect, these results do not replicate across different datasets and model specification (Table 1). Moreover, while the actual effects fall in the range between 1.3% and 4.7% and are therefore relatively small, they do reach significance and are driven by our economic-games stimuli.

### Reaction Time across Belief Conditions and Task Domains

Figure 2B shows the mean reaction times across Task Domains and Belief conditions. We analyzed the log reaction times of correct trials only, and found significant main effects of Belief (X^2^(1) = 4.52, *p* = 0.033) and Task Domain (X^2^(1) = 63.70, *p* < 0.001) and a significant interaction between Belief and Task Domain (X^2^(1) = 69.51, *p* < 0.001). Follow-up pairwise tests of the interaction were conducted using the free method from the multcomp package via the Sidak correction. Results indicate a significant difference between economic games and life stimuli in the Outcome conditions (estimate = −0.469, t = −18.81, *p* < 0.001), while no significant difference between the economic games and life stimuli was observed in the Belief conditions (estimate = −0.046, t = −1.88, *p* = 0.064). These results indicate that in the Outcome condition response times were significantly faster for economic games, while participants spend about equally long answering questions about beliefs in the economic-games and standard FBT. This again agrees with the deviation of behavior with this type of stimulus from the other vignette stimuli. Note that our fMRI models implicitly control for these reaction time differences by implementing a variable-epoch model for all question period regressors (Grinband et al., 2008).

### FMRI results

#### Mentalizing Effects during Belief Formation in the Vignette Period across task Domains

In our initial analyses, we focus on the vignette period during which participants were required to form a belief about the protagonists’ mental state by reading about a sequence of events. To test whether our economic-game vignettes elicit similar activation patterns in social cognition regions as standard FBT vignettes, we first identify the neural correlates of mentalizing via the contrast belief > outcome, and did this separately for economic and life story vignettes. For the life story vignettes, our results replicate previous findings (Saxe & Kanwisher, 2003, Bruneau et al., 2012; van Overwalle, 2009; Schurz et al., 2014), as we find significant activation in bilateral temporal parietal junction (left TPJ: −51, −55, 29, k=678; right TPJ: 54, −49, 23, k=1094), dorsal medial prefrontal cortex (dmPFC, 0, 47, 32, k=427), precuneus (3, −58, 38, k=145), and also bilateral inferior frontal gyrus (IFG) (left IFG: −30, 20, −19, k=72; right IFG: 57, 26, −10, k=140) (**Figure 3A**, Table 3). For the novel economic game vignettes, we find a less distributed set of social cognition regions that include dmPFC (−9, −53, 29, k=246), left TPJ (−54, −70, 32, k=106), right temporal pole (51, −10, −37, k = 310), and left temporal gyrus / temporal pole (−48, −1, −25, k=276) (**Figure 3B**, Table 3).

**Figure 3.**
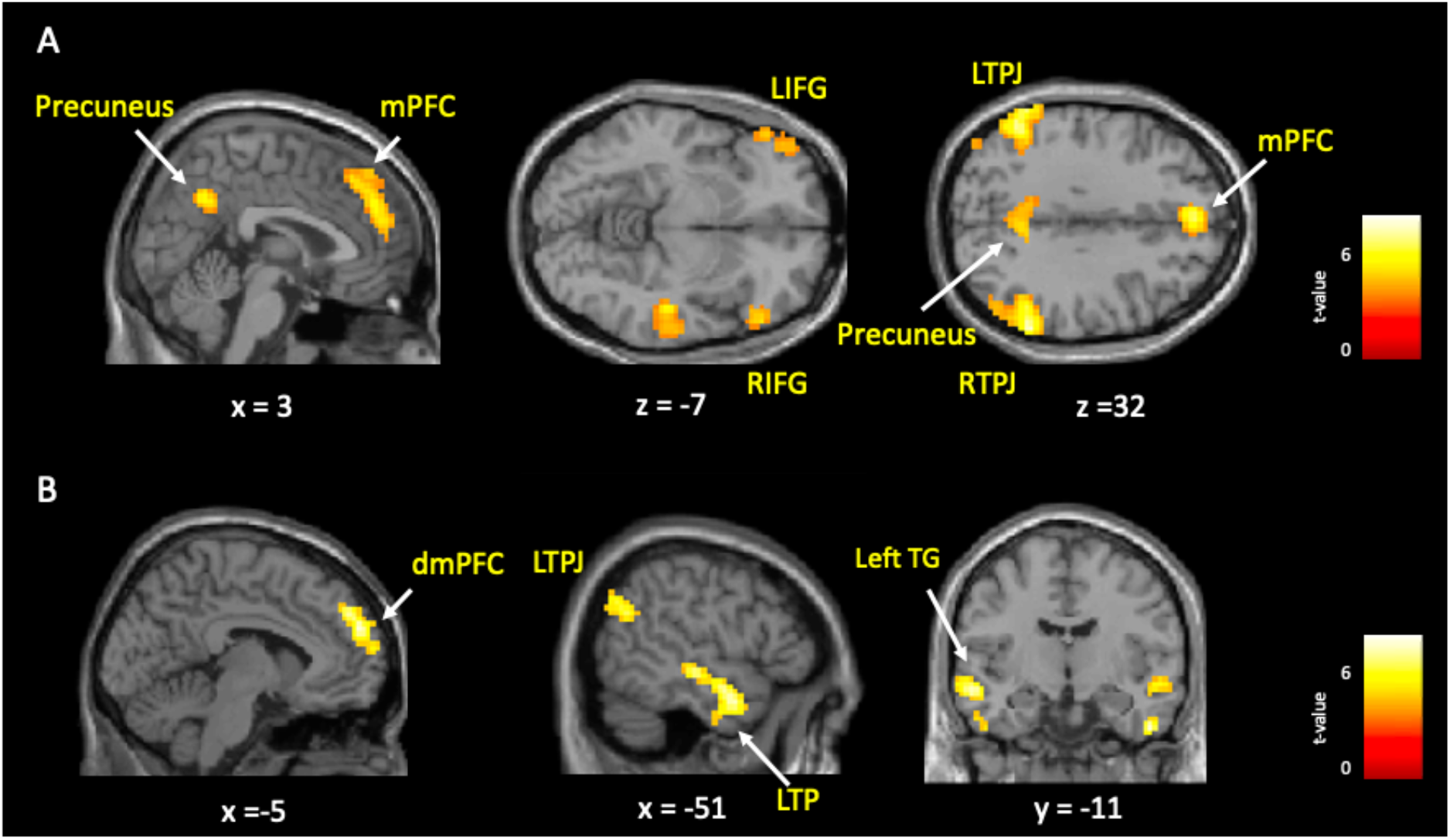
Whole brain analysis of belief activations during the vignette period for the contrast belief > outcome in the life story domain (A) and economic game domain (B). Results show consistent activations in theory of mind regions in both tasks, particularly in dmPFC and left TPJ. Results shown here were FWE-corrected at cluster level with a cluster-forming threshold of *p* < 0.001.

**Table 3.** Whole brain analysis of mentalizing effect during vignette period in the life story and economic game domain (*p* < 0.05 FWE corrected at cluster-level).

| Structure | L/R | Cluster Size | x | y | z | Peak t |
| --- | --- | --- | --- | --- | --- | --- |
| *Belief > Outcome (Life story)* |  |  |  |  |  |  |
| TPJ | R | 1094 | 54 | -49 | 23 | 7.45 |
| TPJ / supramarginal gyrus | L | 678 | -51 | -55 | 29 | 6.99 |
| dmPFC | Bil. | 427 | 0 | 47 | 32 | 5.93 |
| Inferior frontal gyrus | L | 72 | -30 | 20 | -19 | 5.33 |
| Precuneus | Bil. | 145 | 3 | -58 | 38 | 5.29 |
| Inferior frontal gyrus | R | 140 | 57 | 26 | -10 | 5.11 |
| Frontal eye fields | R | 73 | -48 | 20 | 44 | 4.78 |
| Frontal lobe | L | 79 | -57 | 35 | -4 | 4.49 |
| *Belief > Outcome (Economic game)* |  |  |  |  |  |  |
| Middle temporal gyrus / temporal pole | L | 276 | -48 | -1 | -25 | 6.26 |
| dmPFC / superior frontal gyrus | L | 246 | -9 | 53 | 29 | 5.84 |
| Temporal pole | R | 310 | 51 | -10 | -37 | 5.78 |
| TPJ / angular gyrus | L | 106 | -54 | -70 | 32 | 5.05 |

To test the overlap of these two networks, we performed a conjunction analysis of the FWE-corrected maps shown in Figure 3 reflecting belief activations (belief vs. outcome) in the economic-game and story-based task domains. Thereby we examine which voxels showed significant belief-based activation across both versions of the false-belief task, i.e., the life story and economic games domain. The conjunction analysis identified significant overlap in social cognition regions for both domains, specifically in left TPJ (−51, −61, 26, k=91), dmPFC (−6, 47, 35, k=46), right temporal gyrus (48, −25, −4, k= 158) (**see Figure 4)**. Moreover, we extracted activation patterns from regions that showed significant activation in both the life story and economic game conditions and plot their time course. Inlets in Figure 4 illustrate that, in both the life story and economic game vignettes, in accordance with the relatively sustained nature of this task phase activity in these regions rises after about 5 seconds and, importantly, shows higher peak values in the belief compared to outcome conditions. These results support the notion that this network of regions is involved in mentalizing in both domains, namely life stories and economic games.

**Figure 4.**
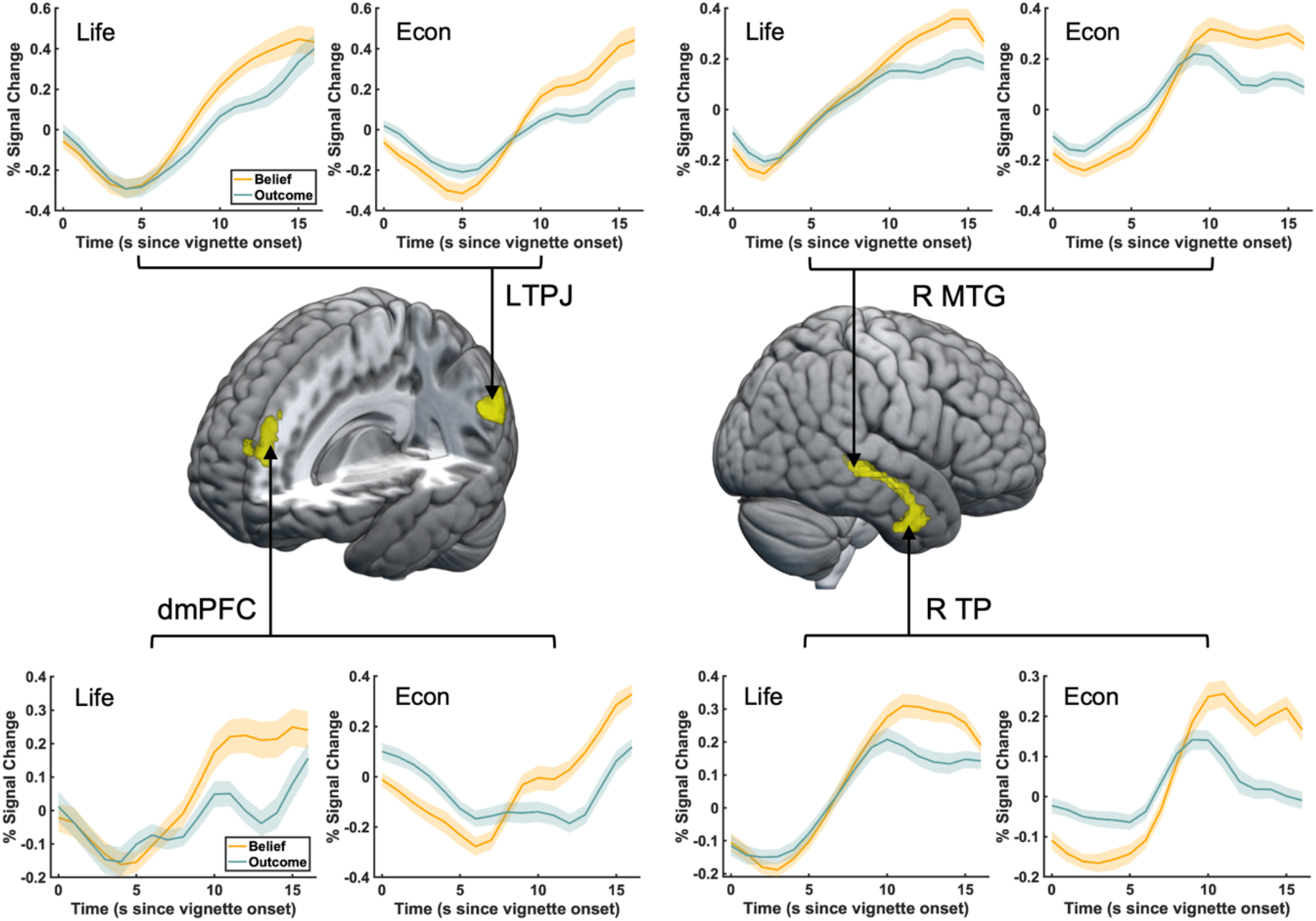
Conjunction analysis during vignette period. A conjunction analysis showed significant overlap between the economic-games and standard FBT in a wider network of social cognition regions, including left TPJ, dmPFC and right middle temporal gyrus/temporal pole. Inlets show time courses of significant activations plotted separately for the life story and economic game domain. Time courses were extracted from voxels in the regions identified by the conjunction analysis, which were further thresholded to separate clusters in middle temporal gyrus. The shaded area denotes the standard error of the percentage signal change.

#### Mentalizing Effects during Belief Inferences in the Question Period across Task Domains

Next, we investigated the period during which participants answered questions concerning the events described in the vignettes. This period required participants to make inferences about the understanding they formed about the protagonists’ beliefs and intentions from the sequence of events described in the life stories and economic interactions to correctly answer the incentivized questions. Since this period required an integration of the information gathered during the vignette period with what was asked in the question, we expected more extended activation patterns that primarily include social cognition regions during this period. We again contrasted belief vs. outcome conditions to test the effect of mentalizing and did so separately for economic game and life story vignettes. In the life story domain, shown in **Figure 5A** and **Table 4**, we identified three large clusters with peaks in precuneus (−3, −67, 32, k=13923), left temporal pole (extending into TPJ; −54, −4, −34, k=219), and left dlPFC (extending into dmPFC; −24, 44, 35, k=524). For questions concerning economic games, shown in **Figure 5B** and **Table 4**, we identified a network that includes bilateral temporal gyrus, with the left region extending into TPJ (−57, −28, −1, k=2698), right temporal pole (45, 8, −28, k=976), as well as dmPFC (−9, 59, 32, k=831), right sensorimotor cortex (45, −25, 65, k=131), right posterior cerebellum (24, −73, −37, k=76), right inferior frontal gyrus (51, 26, 2, k=83); as well as right insula (39, y-16, 17, k=119) and right putamen (24, 11, −7, k=120).

**Figure 5.**
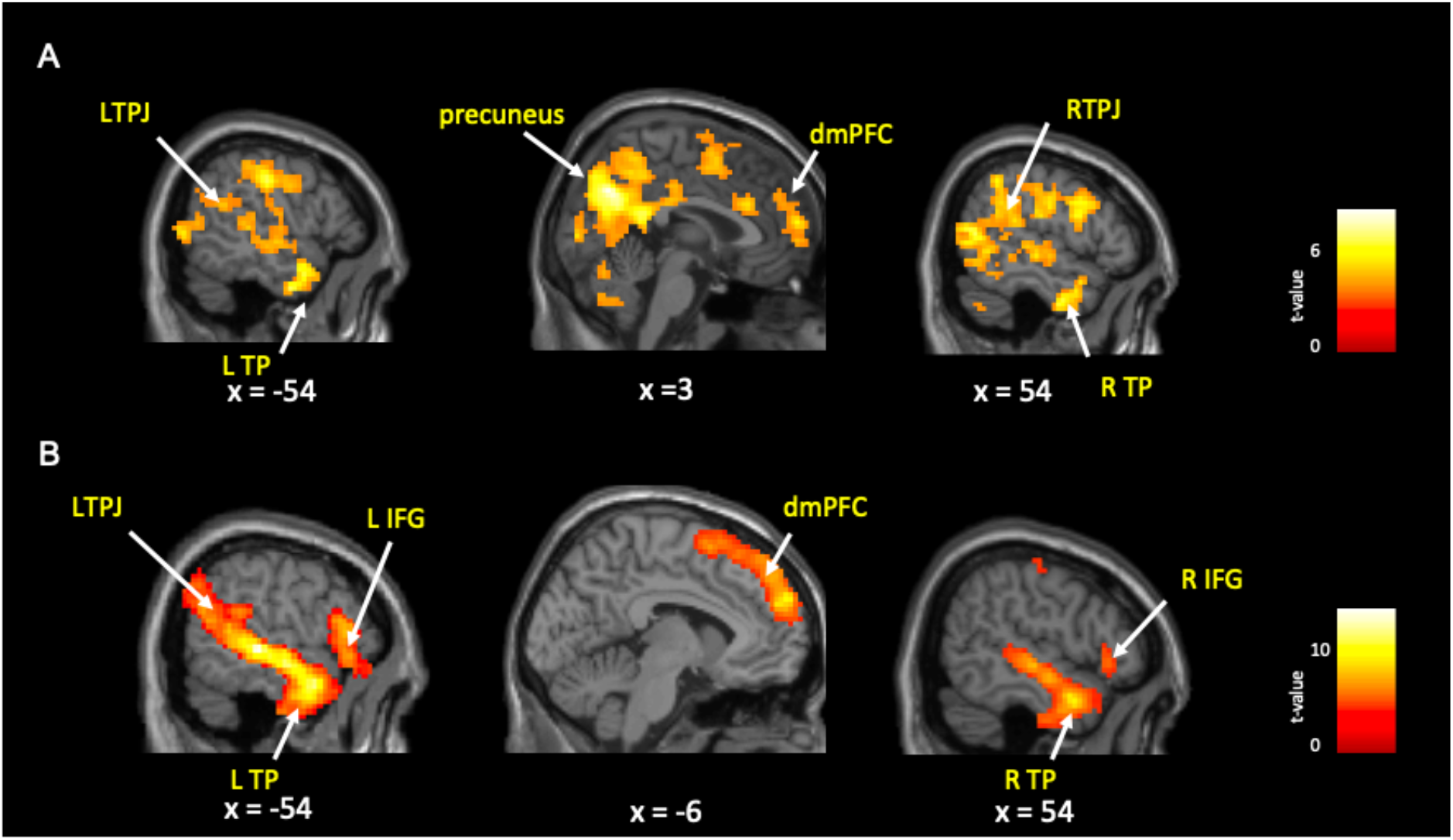
Whole brain analysis of belief activations during the question period for the contrast belief > outcome in the life story domain (A) and economic game domain (B). Results show consistent activations in theory of mind regions in both tasks, particularly in dmPFC, bilateral TPJ and temporal pole. Results shown here were FWE-corrected at cluster level with a cluster-forming threshold of *p* < 0.001.

**Table 4.** Whole brain analysis of mentalizing effect during question period in each Task domain (*p* < 0.05 FWE corrected at cluster-level). Regions listed in italics are subclusters within larger activation clusters. Subclusters were further identified using small volume correction (svc) for each TPJ cluster from the neurosynth map obtained via an association test for the term “mentalizing”.

| Structure | L/R | Cluster Size | x | y | z | Peak t |
| --- | --- | --- | --- | --- | --- | --- |
| *Belief > Outcome (Life story)* |  |  |  |  |  |  |
| Precuneus (extending into) | Bil. | 13923 | -3 | -67 | 32 | 7.44 |
| *TPJ (svc)* | *L* | 22 | -57 | -58 | 23 | 4.33 |
| *TPJ (svc)* | *R* | 146 | 48 | -67 | 14 | 6.1 |
| Temporal pole | L | 219 | -54 | -4 | -34 | 5.75 |
| DLPFC | L | 524 | -24 | 44 | 35 | 5.39 |
| *Belief > Outcome (Economic games)* |  |  |  |  |  |  |
| Superior temporal gyrus (extending into) |  | 2698 | -57 | -28 | -1 | 12.67 |
| *TPJ (svc)* | *L* | 164 | -63 | -61 | 20 | 7.41 |
| Temporal pole | R | 976 | 45 | 8 | -28 | 8.77 |
| Medial PFC | Bil. | 831 | -9 | 59 | 32 | 8.63 |
| Precentral gyrus | R | 112 | 66 | -4 | 29 | 5.72 |
| Posterior insula | R | 119 | 39 | -16 | 17 | 5.70 |
| Sensorimotor cortex | R | 131 | 45 | -25 | 65 | 5.70 |
| Posterior cerebellum | R | 76 | 24 | -73 | -37 | 5.68 |
| Inferior frontal regions | R | 83 | 51 | 26 | 2 | 5.30 |
| Putamen | R | 120 | 24 | 11 | -7 | 4.91 |

**Table 5.** Results from conjunction analyses for vignette and question periods. Regions activated in both the economic game and life story domain were identified by conjoining the two statistical maps, which were each thresholded via a cluster-forming *p* value of *p* < 0.001 and an FWE-corrected cluster threshold. Additional regions are listed in italics that reflect a less conservative conjunction analysis based only on a cluster-forming threshold of *p* < 0.001.

| Structure | L/R | Cluster Size | x | y | z |
| --- | --- | --- | --- | --- | --- |
| *Conjunction of mentalizing effect during vignette period* |  |  |  |  |  |
| Superior temporal gyrus | R | 158 | 48 | -25 | -4 |
| dmPFC | Bil. | 46 | -6 | 47 | 35 |
| TPJ | L | 91 | -51 | -61 | 26 |
| *TPJ* | R | 18 | 48 | -55 | 23 |
| *Middle temporal gyrus* | L | 11 | -60 | -10 | -13 |
| *Temporal pole* | L | 10 | -54 | -7 | -31 |
| *Precuneus* | Bil. | 7 | -6 | -55 | 29 |
| *Conjunction of mentalizing effect during question period* |  |  |  |  |  |
| Temporal pole | L | 203 | -54 | 5 | -25 |
| Temporal pole | R | 434 | 48 | -7 | -37 |
| Middle temporal gyrus / TPJ | L | 455 | -54 | -28 | -4 |
| Cerebellum | R | 39 | 24 | -73 | -37 |
| Putamen | R | 43 | 24 | 17 | -7 |
| dmPFC | Bil. | 55 | -6 | 56 | 26 |
| SMA / pre-SMA | Bil. | 44 | -3 | 8 | 65 |
| *Cerebellum* | L | 25 | -27 | -76 | -40 |
| *Precuneus* | Bil | 18 | -3 | -55 | 29 |
| *Temporal pole* | L | 5 | -27 | 8 | -31 |
| *Pre-motor area* | L | 5 | -48 | -4 | 50 |

Next, similar to the approach for the vignette reading period, we examined the overlap of the networks recruited in both the life story and economic game domains via a conjunction analysis of the FWE-corrected maps shown in Figure 5 reflecting belief activations (belief vs. outcome) in the economic-game and story-based task domain. The conjunction results are shown in **Figure 6** and confirm that significant belief-based activation occurred in a network of overlapping regions in the life story and economic game domains. Areas that are activated across these conditions include the dmPFC (−6, 56, 26, k = 55), left middle temporal gyrus extending into TPJ (−54, −28, −4, k = 455), left temporal pole (−54, 5, −25, k = 203), supplementary motor cortex (−3, 8, 65, k = 44), right temporal gyrus extending into temporal pole (48, −7, −37, k = 434) and right posterior cerebellum (24, −73, −37, k = 39).

**Figure 6.**
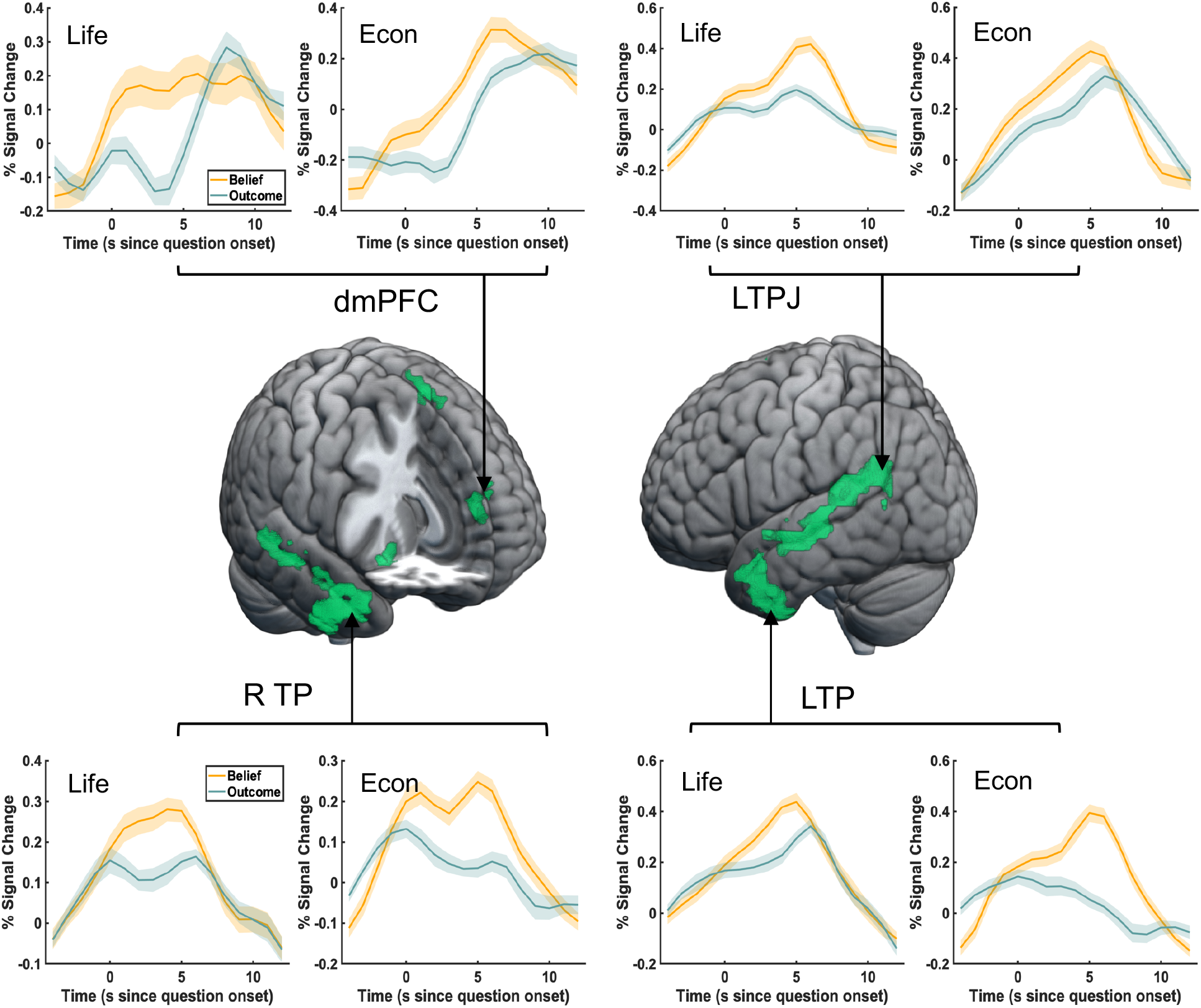
Conjunction analysis during question period. A conjunction analysis showed significant overlap in a wider network of mentalizing regions, including left TPJ, dmPFC and bilateral temporal pole. Inlets show time courses of significant activations plotted separately for the life story and economic game domain. Time courses were extracted from voxels in the regions identified by the conjunction analysis, which were further thresholded to separate clusters in middle temporal gyrus. The shaded area denotes the standard error of the percentage signal change.

**Figure 7.**
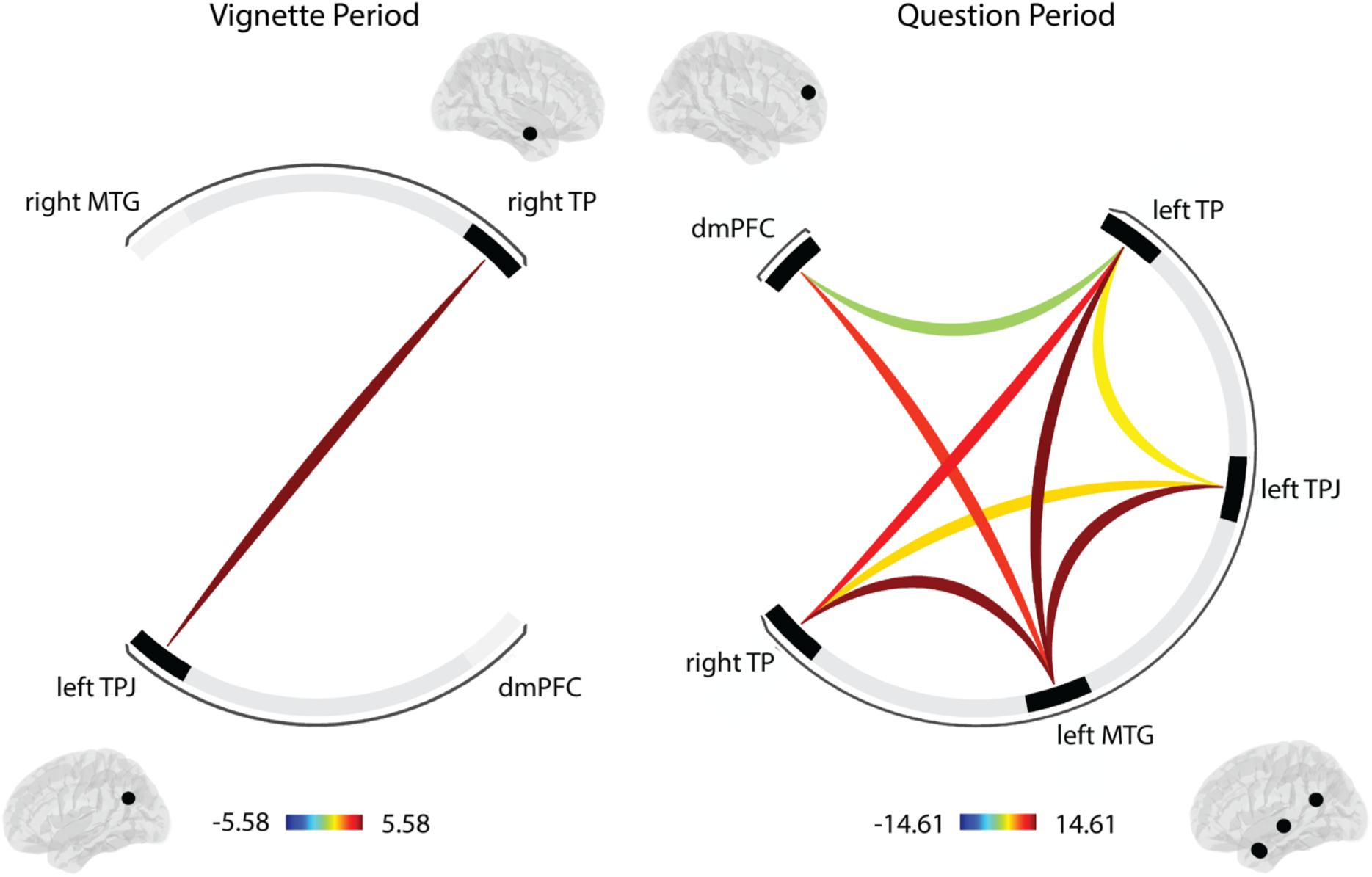
Functional connectivity among ROIs during the vignette and question periods. ROI-to-ROI analyses show heightened connectivity during belief relative to outcome conditions between left TPJ and right TP during the vignette period (left ROI-ring display, TFCE = 5.58, FWE-corrected *p* = 0.044) and extensive interconnectivity among ROIs during the question period (right ROI-ring display, TFCE = 14.91, FWE-corrected *p* = 0.009).

Moreover, we extracted activation patterns from regions that showed significant activation in both the life story and economic game conditions and plotted the respective time courses. Inlets in **Figure 6** illustrate that, in accordance with the more transient nature of this task phase, activity in these regions rises almost immediately after the onset of the question period and peaks at about 6 seconds. Time courses also show a larger peak in the belief compared to outcome conditions for both the life story and economic game vignettes. These results support the notion that this network of regions is involved in mentalizing in both the life stories and economic game domains during the question period.

#### Functional connectivity during mentalizing

In our final analyses, we asked the question to what extent the regions identified by the conjunction analyses between our economic game and story-based vignettes are functionally interconnected with other social cognition regions during mentalizing. To this end, we conducted generalized Psychophysiological Interaction (gPPI) analyses. First, using ROI-to-ROI analyses, we inspected the belief-based (belief vs. outcome) interconnectivity within our set of ROIs during each of the task phases. Next, using seed-based whole-brain analyses, we assessed whether additional target regions showed stronger positive connectivity with our seed regions during belief relative to outcome conditions. Analyses were conducted separately for the vignette and question periods, and for each ROI-to-ROI and seed-based analysis, we used as seeds those regions that were identified by the conjunction analysis for that specific period (see methods).

During the vignette period, we find that the left TPJ shows significant interconnectivity with right TP (TFCE = 5.58, FWE-corrected *p* = 0.044), indicating relatively restricted interconnectivity within our network of ROIs. This could be due to a mismatch between the sustained nature of the vignette period and how regions in fact communicate throughout this period, such that the fluctuation of transient and repeated communication between regions might not be picked up by the current regression analysis. In the shorter question period, we see extensive interconnectivity between all the social cognition regions we included as ROIs (TFCE = 14.91, FWE-corrected *p* = 0.009). This indicates that preparing an answer that involves an understanding of beliefs requires strong cross-talk between social cognition regions.

For our whole-brain gPPI analyses, we observe an interesting pattern that highlights the role of right TPJ during the vignette period, which is a target region of both the left TPJ (left to right TPJ: 58, −52, 30, k = 126, cluster-level FWE-corrected *p* = 0.0253), and the right TP (right TP to right TPJ: 52, −54, 26, k = 570, cluster-level FWE-corrected *p* < 0.0001) during belief relative to outcome vignettes (**Figure 8)**. This result is interesting, as it confirms the role of the right TPJ in mentalizing, which we do not find in conjunction analyses reported above, and shows the importance of a wider interconnected set of regions involved in mentalizing during the vignette period. We also find reduced connectivity between the TP seed region and a target in sensorimotor area (−26, −30, 66, k = 177, cluster-level FWE-corrected *p* = 0.0038).

**Figure 8.**
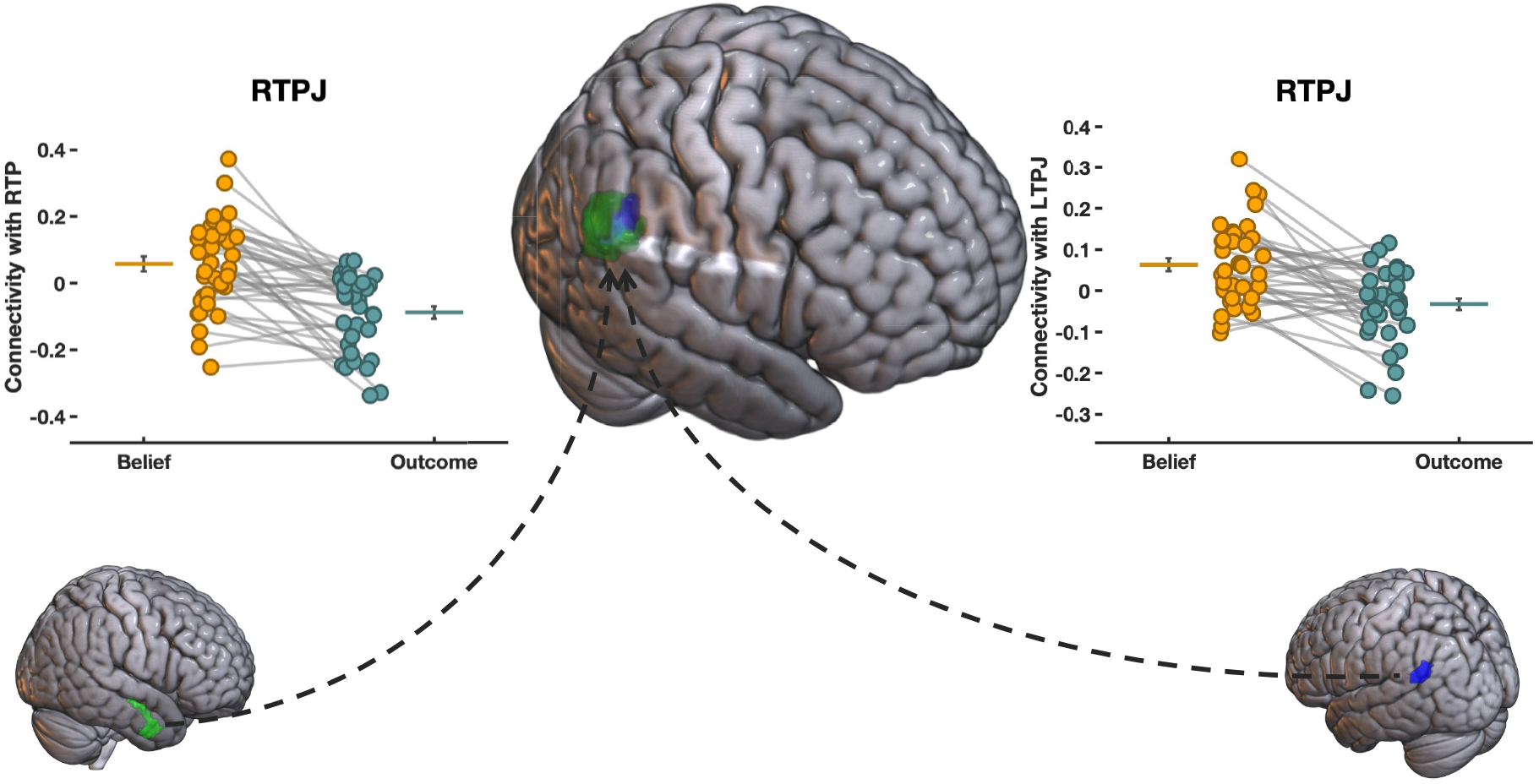
Whole-brain gPPI analysis of belief-based effective connectivity during the vignette period. gPPI analyses show heightened connectivity during belief relative to outcome conditions between left TPJ seed and right TPJ target during the vignette period (58, −52, 30, k = 126, clusterlevel FWE-corrected *p* = 0.0253) as well as between right TP seed and right TPJ target (52, −54, 26, k = 570, cluster-level FWE-corrected *p* < 0.0001).

During the question period, we find enhanced belief-based connectivity between the left TPJ and its target in right cerebellum (Figure S2; 24, −78, −18, k = 169, cluster-level FWE-corrected *p* =0.0059). Finally, the dmPFC shows enhanced belief-based connectivity with a region in superior parietal lobe that extends to precuneus (Figure S3; −24, −66, 48, k = 141, cluster-level FWE-corrected *p* = 0.0150).

## Discussion

An important question in the field of social neuroeconomics is whether the activations within brain regions that are meta-analytically associated with mentalizing and that are also consistently involved in decisions in the context of interactive economic games (e.g., Alos-Ferrer and Farolfi, 2019; Fehr and Camerer, 2007; Engelmann et al, 2019; Rilling and Sanfey, 2011) indeed reflect mentalizing about interaction partners. While this conjecture is theoretically plausible and is supported by the stark overlap of activation patterns across a variety of tasks that are associated with belief inferences (Mar, 2011; Molenberghs et al., 2016; Schurz et al., 2014; Van Overwalle, 2009), it is important to directly compare and identify the overlap between the neural systems engaged in mentalizing across different contexts, including in life events but also in an economic games context, in the same participants using the same task. The goal of the current study was therefore to move beyond reverse inference and address this gap in the literature using a novel version of the false belief task that required our participants to make belief-based inferences in the context of economic game scenarios.

Our fMRI results indeed identify strong overlap between the networks engaged during the standard false-belief task and a modified version that requires an understanding of economic games to correctly infer beliefs of interaction partners in hypothetical economic games. This shows that our novel economic-games false-belief task, which asked participants to observe two agents interact in the trust and ultimatum game and make inferences about their beliefs, reliably activated canonical social cognition regions. Specifically, using conjunction analyses we find two regions that show enhanced activity during belief-based (relative to outcome-based) inferences during both variants of the task, namely the left TPJ and dmPFC. This finding is in line with a series of previous meta-analyses on the neural underpinnings of mentalizing, which consistently pinpointed these two nodes as core areas for mentalizing across different paradigms, including economic games (Mar, 2011; Molenberghs et al., 2016; Schurz et al., 2014; Van Overwalle, 2009). Moreover, we find that these regions are involved in reasoning about others’ beliefs during two periods of our task: the vignette period, during which participants need to read and understand the beliefs of others, and the question period, which required them to integrate the information gathered via the vignettes and answer a brief question about the protagonists’ beliefs. The consistency of the activation overlap across the different task types and task periods further underlines the importance of these regions for belief-based inferences. Moreover, these results underline the importance of TPJ and dmPFC for belief-based inferences in the domain of economic games. Jointly, our results substantiate the notion that the commonly observed activation of social cognition regions during interactive economic games, particularly the TPJ and dmPFC, reflects mentalizing (Rilling and Sanfey, 2011; Engelmann et al., 2019; Fehr and Camerer, 2007).

The important role of the temporoparietal junction in mentalizing is further underlined by effective connectivity analyses. During the vignette period, the left TPJ shows enhanced belief-based connectivity with right TPJ, and right TPJ is a target of right temporal pole (Figure 8). This shows that even if the TPJ does not show bilateral activation in conjunction analyses, effective connectivity patterns implicate bilateral TPJ during mentalizing in the vignette period. Moreover, connectivity patterns also underline the importance of cross-talk within a wider network of social cognition regions that include bilateral TPJ, bilateral TP and dmPFC, when participants make belief-based inferences that involve mentalizing during the question period.

Our results furthermore indicate that there is a more extensive network of regions that are involved in belief-based inferences across the two task versions. This is clear from two types of analyses: 1) Conjunction analyses of the overlap of activation patterns across standard and economic-game false-belief task versions, and 2) effective connectivity analyses involving the regions identified in these conjunction analyses. The conjunction analyses identified more extended belief-based activation in right temporal pole (extending into right middle temporal gyrus) during the vignette period, and bilateral temporal pole during the question period. Moreover, the TP also showed heightened belief-based connectivity with target regions associated with social cognition, including the right TPJ during the vignette period (Figure 8), and left TPJ and left MTG during the question period. Our results of heightened belief-based activity and connectivity of the temporal pole agree with its roles in semantic memory, face recognition, and theory of mind (Gainotti et al., 2003; Gentileschi et al., 2001; Olson et al., 2007), as all of these are social cognitive skills that support belief-based inferences (e.g., Patterson et al., 2007). Moreover, this result is consistent with previous studies on the neural correlates of social cognitive (Frith & Frith, 2006) and social affective mechanisms (Völlm et al., 2006).

As part of a more extended network of social cognition regions involved in mentalizing, the cerebellum deserves some additional discussion. Specifically, we find significant activation in right posterior cerebellum during belief-based inferences in the question period (Table 4), and furthermore, the right posterior cerebellum is found as a target of left TPJ in connectivity analyses. Our results therefore substantiate the importance of the cerebellum as a region that supports mentalizing in important ways, but that falls outside of the typical social cognition areas within cerebral cortex. In fact, a recent meta-analysis based on 350 fMRI studies provides strong support for the notion that the cerebellum subserves important social cognitive functions, particularly when a certain level of abstraction is required (Van Overwalle et al., 2014). These social cognitive functions include mirroring others’ behavior, mentalizing, and the representation of abstract concepts in social contexts (e.g., group stereotypes). Our fMRI results support the hypothesis that the cerebellum is involved in belief-based inferences about others.

Moreover, the location of the cerebellum activation found in the current study corresponds well with what has been reported previously. Van Overwalle et al. (2014) suggest that right hemisphere lateralization of cerebellum was specifically associated with mentalizing tasks that require language processing (Stoodley & Schmahmann, 2009), which matches the results reported here. Van Overwalle & Mariën (2016) examined the functional connectivity between cerebellum and cerebrum for mentalizing across five studies with high level of abstractness (e.g., judgement of others’ traits, group stereotypes). They found significantly higher functional connectivity between right posterior cerebellum and bilateral TPJ and dmPFC. Our results partially validate this prior finding, showing significantly higher belief-based functional connectivity between left TPJ and right posterior cerebellum during the question period. Taken together, our fMRI results are consistent with previous findings implicating the cerebellum in social cognitive processes, and lend further support to the notion that the cerebellum is involved in belief-based inferences about others. It is therefore important for future studies in social neuroscience and social neuroeconomics to also examine the results in cerebellum carefully.

### Limitations

As with every experiment, there are a number of limitations that need to be considered. The current paper presents a reanalysis of data from a larger experiment on the effects of anxiety on theory of mind. One of the limitations therefore is that participants completed the task in the context of threat blocks, in which they could experience electric shocks at unpredictable time points, and safe blocks, during which they were free from the threat of electric shocks. This approach is known to induce affective states of anxiety during threat blocks and relative safety during safe blocks (e.g., Engelmann et al., 2015, 2019) and these affective states might enhance or depress the belief-based activation and connectivity of the regions reported in the current paper. We tackle this limitation by controlling for these effects and including the factor threat, as well as each electrical shock moment as regressors of no interest in all of our analyses. Given that these factors should mostly increase noise in our data and work against our results, in conjunction with our activation and connectivity patterns being highly consistent with those previously reported in experiments and meta-analyses of the neural correlates of mentalizing (Mar, 2011; Molenberghs et al., 2016; Schurz et al., 2014; Van Overwalle, 2009), we are confident in the validity of our results.

A second limitation concerns our analyses of two separate periods of the task, the vignette period, during which participants were reading and forming an understanding of the events outlined in the vignette, and the question period, during which participants were asked to make inferences about what they just read. Our experimental design did not include jitter between these two periods, which would have allowed us to better separate the hemodynamic response across vignette and question periods. We made this decision for three reasons: 1) To allow better comparison with previous studies (e.g., Saxe & Kanwisher, 2003; Young et al., 2010a; Young et al., 2010b), 2) to ease the cognitive burden on our participants that jitter might have imposed, as suggested by results from our behavioral pilot study and 3) to keep the experiment relatively short. Moreover, this limitation is qualified by the BOLD patterns shown in Figures 4 and 6. We find during both task periods that BOLD responses follow the expected pattern given the cognitive demands of that period. During the vignette period BOLD responses rise to peak between 10 and 15 seconds, reflecting the more sustained nature of social cognitive processes required to understand a sequence of events during this period. During the question period, we observe that the BOLD response starts from a low activation level (around zero percent signal change) and rises to peak at around 5 seconds, reflecting the more transient nature of social cognitive processes during this period that is consistent with the average response time of 2.61 seconds during this period. Our findings that the BOLD responses during vignette and question periods follow patterns that are consistent with the cognitive demands of each period, and that they start from a low activation level in the question period in regions that show overlap with those activated during the preceding period (left TPJ and dmPFC), therefore mitigate this concern.

Finally, we need to point out that the control condition in the economic games false-belief task is different from the control condition in the standard false-belief task. While in the standard false-belief task, we used a story-based outcome condition, in the Economic Game-Outcome condition our participants were asked to calculate the payoff of one of the interaction partners based on the rules of the economic game in question (TG or UG). While this leads to somewhat different behavioral results in this condition (Figure 2), we argue that the economic game outcome condition is nonetheless an ideal control condition for belief-based inferences made in the context of economic game vignettes. This is the case because participants need to apply the same understanding of economic games in both the belief and outcome conditions, but focus on different aspects of the social interaction, namely the interaction partners’ beliefs compared to their payouts (which are also a result of the social interaction). Furthermore, including the economic games outcome condition allowed us to ensure that participants understand the rules of the economic games and were able to calculate their payouts across different contexts.

### Conclusions

Our findings lend support to the notion that activations within the social cognition network that have consistently been observed during decisions in the context of interactive economic games reflect mentalizing about interaction partners. We addressed this question here by developing a novel version of the false-belief task that is based on interactions in economic games, specifically the trust game and ultimatum games. Correctly answering questions about the beliefs of the interaction partner in the economic games false-belief task requires an understanding of the rules of these games. Comparing activation patterns during the standard story-based false-belief task with a novel game-theoretic false-belief task in the same participants, we identify overlap between the neural systems engaged in mentalizing. Specifically, our conjunction analyses identify two regions that show enhanced activity during belief-based (relative to outcome-based) inferences during both variants of the task, namely the left TPJ and dmPFC, which is in line with results from previous meta-analyses (Mar, 2011; Molenberghs et al., 2016; Schurz et al., 2014; Van Overwalle, 2009). Moreover, we find an extended network of regions that are important for mentalizing during both task versions, with the temporal pole being prominently represented in conjunction and connectivity analyses, and the right TPJ showing enhanced connectivity with left TPJ and right TP during the vignette period. Jointly, our results support the notion that mentalizing during belief formation and inferences are supported by social cognitive processes in a wider network of social cognition regions that include bilateral TPJ, TP and dmPFC as central nodes. Importantly, this is the case in the context of economic games and standard false-belief tasks.

## Acknowledgements

We are grateful to Alfonso Nieto-Castanon for the helpful and fast assistance with the CONN toolbox. JBE gratefully acknowledges startup funds from the Amsterdam School of Economics that supported this work.

## Notes

https://osf.io/3eg56/?view_only=face48878dd144848d26f1c7d3c47d31

### Competing Interest Statement

The authors have declared no competing interest.

### Summary of Updates

Additional control analyses were added

