## Supplement for "Mentalizing in an economic games context is associated with enhanced activation and connectivity in left temporoparietal junction"

For

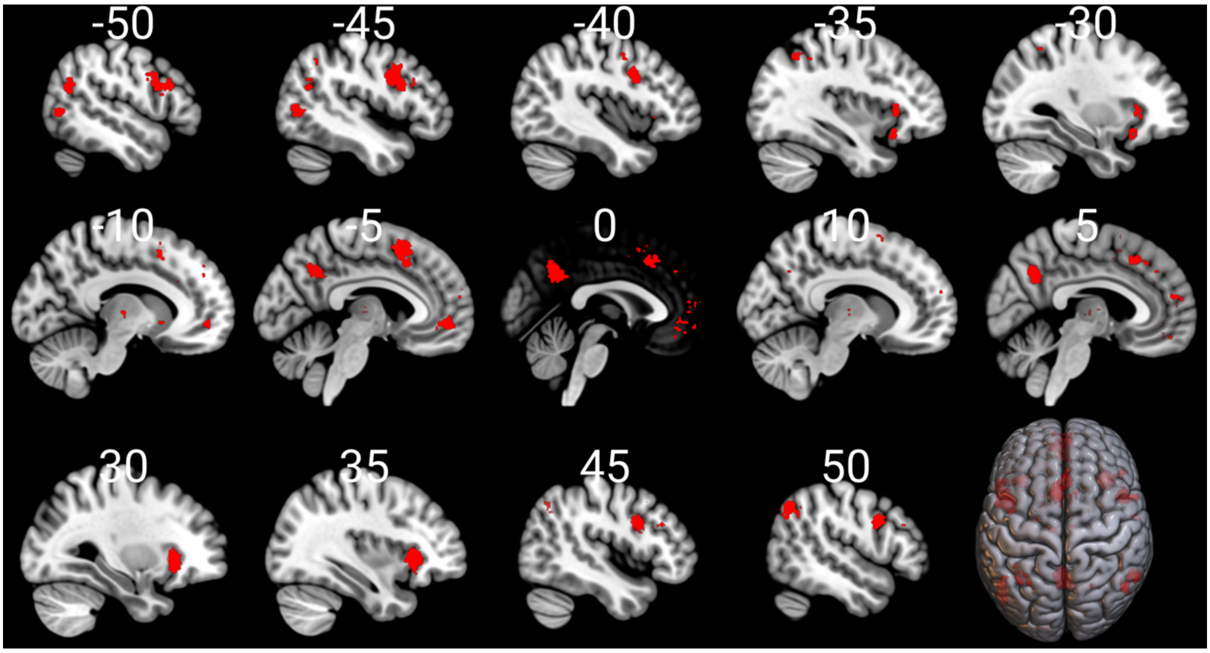


**Figure S1.** A conjunction analysis of two Neurosynth meta-analyses for the terms "game" (N=176) and "mentalizing" (N = 151) reveals overlap across these two tasks in canonical social cognition regions including bilateral TPJ, precuneus/PCC, and multiple clusters within dmPFC. Additional regions that show overlap include the vmPFC, AI and dlPFC. Note that uniformity tests were used for both meta-analyses to create this conjunction map due to a lack of results for the association test for "game". The conjunction was computed by multiplying the two neurosynth meta-analyses maps using the SPM tool imcalc.

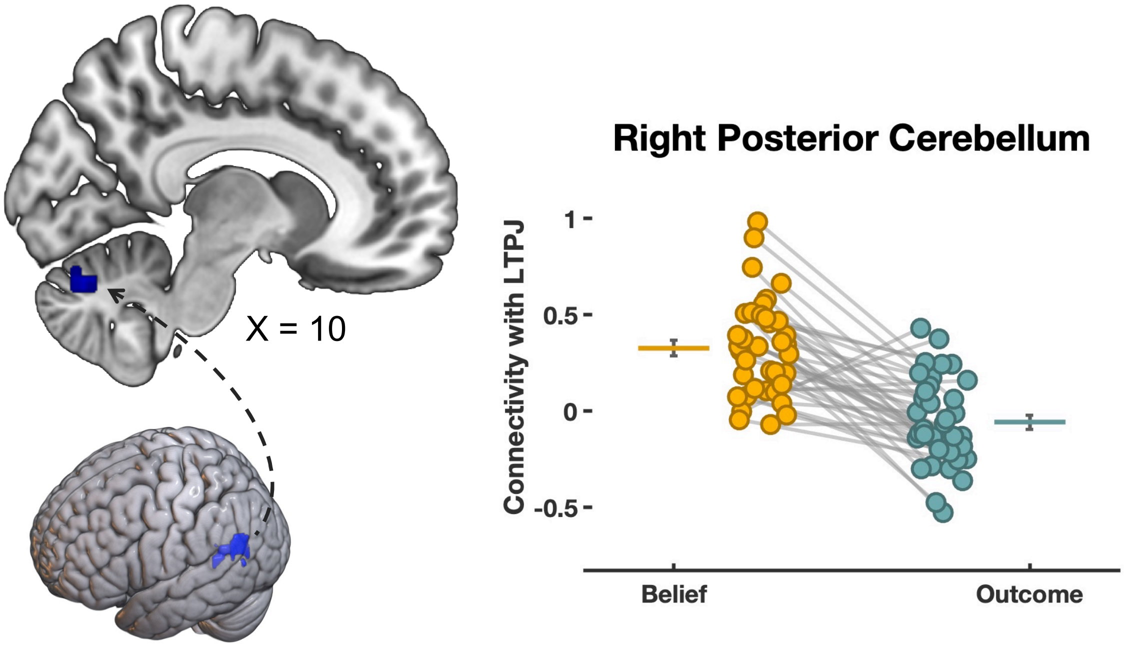


**Figure S2.** Question period whole-brain gPPI results showing the belief-based connectivity increases between the left TPJ seed and right posterior cerebellum target (24, -78, -18, k = 169, cluster-level FWE-corrected $p$ =0.0059).

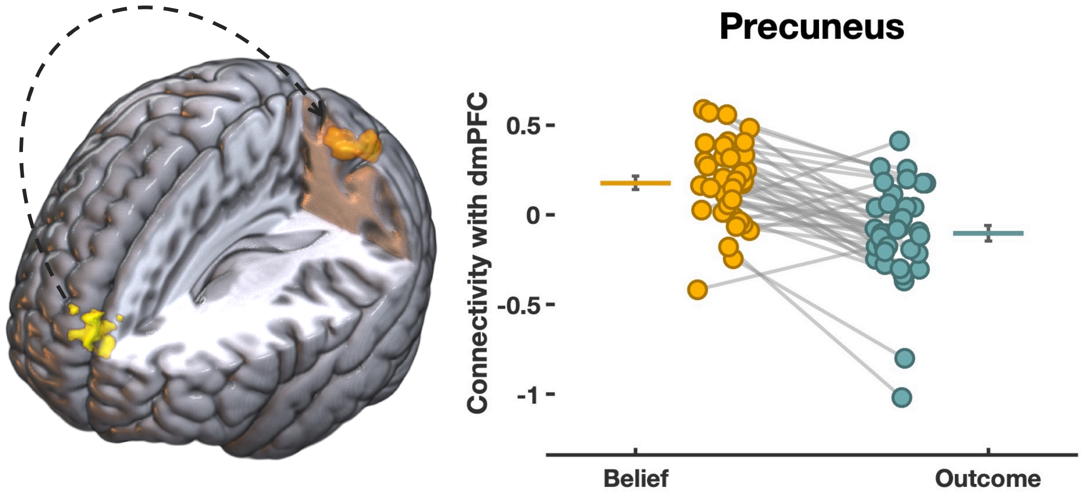


**Figure S3.** Question period whole-brain gPPI results showing the belief-based connectivity increases between the dmPFC seed and target in Precuneus/superior parietal lobe (-24, -66, 48, k = 141, cluster-level FWE-corrected $p$ = 0.0150).

**Manipulation Check: The effect of strategy use on behavioral and imaging results**

We assessed whether participants used a strategy in the exit questionnaire used after completion of the fMRI experiment. Specifically, we asked participants whether they used a specific strategy to answer the questions. If they answered yes, we asked them in an open-ended question to describe the strategy they used. A number of participants (43%) indeed reported using some strategy when answering the questions. The most common strategy that participants reported indicates that they scanned for particular words in the vignette, such as rejected or accepted. We re-analyzed our imaging and behavioral results by adding a binary "strategy" covariate to our models. We pursued two goals in these control analyses: 1. We checked whether controlling for the strategy by including a binary covariate for strategy in our imaging and behavioral models changes the results; and 2. We checked whether there are differences across those participants that report using a strategy and those that do not at the behavioral and imaging level. Controlling for strategy does not change results at the behavioral and imaging level, with all (non-) significant effects remaining at the behavioral level, and slight changes in the extent of the activation patterns at the imaging level (see tables S1 and S2, imaging tables are largely equivalent to the ones already reported in the paper and therefore not reported here, but are available upon request).

Additionally, we probed for behavioral and activation differences as a function of strategy. We find that at the behavioral level there are no significant interactions with our treatments for both dependent variables (accuracy and reactions time). More specifically, using a strategy did not impact the accuracy of the answers, as there were not main and interaction effects of strategy (Table S1). Reaction times were marginally faster overall for people reporting having used a strategy, but this did not impact the performance differentially across the two task domains as there were no interaction effects with strategy (life and econ). Moreover, we find no effect of strategy in our neuroimaging results of the economic-games FBT task. In other words, we did not observe differential activations during belief relative to outcome for those subjects that reported using a strategy compared to those that did not. Interestingly, however, neuroimaging results in the standard false belief task were impacted by self-reported strategy use, indicating that strategies affect the neural correlates of the original FBT in the Putamen, anterior cingulate cortex (ACC) and regions within the insula (see Table S3).

| | Strategy control | | | | Strategy modulation | | | |
|---|---|---|---|---|---|---|---|---|
| | Chisq | Df | Pr(>Chisq) | | Chisq | Df | Pr(>Chisq) | |
| Belief | 17.18 | 1 | <0.001 | *** | 16.43 | 1 | <0.001 | *** |
| Life | 1.86 | 1 | 0.17 | | 1.49 | 1 | 0.22 | |
| Threat | 2.36 | 1 | 0.12 | | 2.38 | 1 | 0.12 | |
| Strategy | 0.02 | 1 | 0.90 | | 0.01 | 1 | 0.92 | |
| Belief x Life | 18.31 | 1 | <0.001 | *** | 18.49 | 1 | <0.001 | *** |
| Belief x Strategy | | | | | 0.03 | 1 | 0.85 | |
| Life x Strategy | | | | | 1.36 | 1 | 0.24 | |
| Belief x Life X Strategy | | | | | 0.10 | 1 | 0.76 | |

**Table S1.** ANOVA tables reflecting accuracy results from mixed models including a Strategy control variable in the left columns, and strategy as a moderator variable on the right.

| | Strategy control | | | | Strategy modulation | | | |
|---|---|---|---|---|---|---|---|---|
| | Chisq | Df | Pr(>Chisq) | | Chisq | Df | Pr(>Chisq) | |
| Belief | 4.53 | 1 | 0.03 | * | 4.25 | 1 | 0.04 | * |
| Life | 64.05 | 1 | <0.001 | *** | 63.49 | 1 | <0.001 | *** |
| Threat | 0.49 | 1 | 0.48 | | 0.49 | 1 | 0.48 | |
| Strategy | 3.66 | 1 | 0.06 | | 3.67 | 1 | 0.06 | |
| Belief x Life | 276.86 | 1 | <0.001 | *** | 275.95 | 1 | <0.001 | *** |
| Belief x Strategy | | | | | 0.20 | 1 | 0.65 | |
| Life x Strategy | | | | | 0.00 | 1 | 0.95 | |
| Belief x Life X Strategy | | | | | 0.89 | 1 | 0.35 | |

**Table S2.** ANOVA tables reflecting reaction time results from mixed models including a Strategy control variable in the left columns, and strategy as a moderator variable in the right columns.

| Structure | L/R | Cluster Size | x | y | z | Peak t |
|---|---|---|---|---|---|---|
| *Economic Game* | | | | | | |
| *Vignette Period: (Belief > Outcome) * Strategy* | | | | | | |
| No significant effects | | | | | | |
| *Question Period: (Belief > Outcome) * Strategy* | | | | | | |
| No significant effects | | | | | | |
| *Life Story* | | | | | | |
| *Vignette Period: (Belief > Outcome) * Strategy* | | | | | | |
| Anterior insula | L | 81 | -51 | 5 | -7 | 5.06 |
| Posterior insula | L | 74 | -51 | -13 | 14 | 4.43 |
| Middle cingulate cortex | L | 97 | -12 | 8 | 41 | 5.27 |
| *Question Period: (Belief > Outcome) * Strategy* | | | | | | |
| Cerebellum | Bill. | 242 | 3 | -73 | -13 | 4.90 |
| Putamen | R | 109 | 27 | 2 | 8 | 5.13 |
| Putamen | L | 196 | -30 | -16 | 5 | 4.74 |
| Thalamus | R | 86 | 12 | -7 | 8 | 4.44 |
| Anterior Cingulate Cortex | Bil. | 102 | -3 | 35 | 14 | 4.71 |
| DLPFC | R | 78 | 42 | 47 | 20 | 4.48 |
| Middle Frontal Gyrus | R | 80 | 27 | 38 | 41 | 4.84 |
| Supplementary Motor Area | Bil. | 105 | 0 | -7 | 50 | 4.68 |

**Table S3.** Whole-brain analysis of the interaction between Strategy and the mentalizing effect during the economic game and life story FBT respectively ($p < 0.05$ FWE corrected at cluster-level). DLPFC: dorsolateral prefrontal cortex.
